# Morphological integration of the avian beak facilitates evolution along lines of least resistance

**DOI:** 10.64898/2026.05.04.722668

**Authors:** Ricardo C. Ely, Stefan H. Sommer, Christy A. Hipsley

## Abstract

Innovation of the avian beak has facilitated a grand radiation of >11,000 species, with vast morphological disparity suggesting limited developmental constraints on beak diversification. We assess four macroevolutionary ‘currencies’ – integration, disparity, phenotypic evolutionary rates, and ecological specialization – using 3D beak landmarks for 8,627 species mapped to a complete avian supertree with a resolved genomic backbone. We introduce a Gini coefficient-based metric of ecological specialization, measuring evolutionary time spent across trophic niches. Phylogenetic regressions show that lineages with faster phenotypic rates exhibit stronger beak integration (landmark covariation) and more generalised diets, while beak disparity declines with greater trophic specialization. These results suggest that integration facilitates, rather than constrains, phenotypic evolution, by channeling variation along lines of least resistance. Future work should explore modular structure of the bird beak, which arises from multiple genetic and developmental factors.

## INTRODUCTION

Morphological integration describes the strength and pattern of covariance among physical traits, arising from developmental commonalities (Cheverud 1996), functional interactions (Cheverud 1982; Sansalone et al. 2022), genetic pleiotropy or epistasis (Cheverud 1996; Klingenberg 2014). The concept of integration has risen in popularity in recent decades, as it provides a testable source of constraint on phenotypic change. Correlations among traits may limit evolution, by linking suites of characters such that their joint variation is concentrated along few dimensions of trait space (Felice et al. 2018). At the same time, strong integration has been associated with greater disparity (Goswami and Polly 2010a; Randau and Goswami 2017; Felice et al. 2018; Randau et al. 2019) and elevated evolutionary rates (Evans et al. 2021), potentially channelling the evolution of coordinated traits along adaptive optima (Schluter 1996; Goswami and Polly 2010a). Such differences in integration outcomes underscore interest in this framework as a predictor of macroevolutionary dynamics, as well as a means to reconstruct genetic and developmental factors shaping biodiversity in adult and extinct forms (Felice et al. 2018).

Among the most frequently hypothesized drivers of morphological integration are ecological factors (Berg 1960; Armbruster et al. 2004; Pélabon et al. 2011). Ecological demands surrounding resource acquisition, locomotion, and activity patterns impose selective pressures that can shape the evolution of morphological, physiological, behavioral, and developmental traits, often simultaneously (Dumont et al. 2016) . For example, comparative morphometric studies have documented covariation of functionally linked traits related to mastication and sensory systems in mammal skulls (Cheverud 1982; Goswami 2006), feeding mode and locomotion in jaws and fins of reef fishes (Collar et al. 2008), substrate use and girdle elements in macaques (Conaway et al. 2018), and foraging behaviour and brain size in cichlids (Kotrschal et al. 2014). Quantitative and synthetic work shows these patterns arise from underlying genetic, developmental and biomechanical links (Cheverud 1982; Klingenberg 2008), including specific developmental effectors (e.g., *Runx2*) that mechanistically couple form and function (Sears et al. 2007).

Integration studies have employed a diversity of taxonomic systems (Conner et al. 2014), birds being no exception. Most studies of avian morphological integration have focused on covariation of the beak and neurocranium (Klingenberg and Marugán-Lobón 2013; Bright et al. 2016; Felice and Goswami 2018; Xu and Natale 2024) or other skeletal structures (Orkney et al. 2021), while integration of the beak itself has received less attention. The outer bird beak is composed of the distal upper bill formed by paired (often fused) premaxillae, with the maxillae contributing to the posterolateral margins and nasals and frontals forming the dorsal roof and base (O’Connor and Chiappe 2011; Bhullar et al. 2015; Plateau and Foth 2020; Iyyanar et al. 2023). The lower bill is dominated by paired dentary bones fused at the symphysis, with posterior elements (surangular/angular derivatives) reduced or integrated into a lightweight mandible (Hogg 1983; Plateau and Foth 2020; Crane et al. 2025). Developmentally these shifts reflect altered neural-crest migration and proliferation (Helms and Schneider 2003; Schneider and Helms 2003; Fish et al. 2014), heterochronic changes in craniofacial growth zones (Abzhanov et al. 2004; Mallarino et al. 2011; Bhullar et al. 2012), and modified activity of conserved signaling pathways that suppress odontogenesis (Chen et al. 2000; Wang et al. 2017a), promote premaxillary expansion (Bhullar et al. 2015), and enable formation of the rhamphotheca, producing a kinetic keratin-covered rostrum.

Since their divergence from non-avian therapod dinosaurs in the late Jurassic, bird beaks have diversified into an extraordinary range of morphologies to meet functional demands related to diet (Schluter and Grant 1984; Grant and Grant 2006), thermoregulation (Greenberg et al. 2012), song production (Derryberry et al. 2012), nesting (Sheard et al. 2023), and preening (Clayton et al. 2005). They experienced elevated rates of evolution and high disparity shortly after the KPg extinction event (Cooney et al. 2017), resulting in maximum morphospace packing relatively early in Cenozoic neornithine (crown clade) evolution. This pattern of diversification implies rapid niche differentiation – a fundamental component of adaptive radiations as species specialize by partitioning broader ecological spaces into finer subdivisions (Harmon et al. 2010). Strong covariance of diet and beak shape has been found at shallow phylogenetic depths and island radiations, particularly in Darwin’s finches (Grant and Grant 2006; Foster et al. 2008). However, only weak associations between beak shape and diet have been recovered across broader macroevolutionary scales (Bright et al. 2016; Felice et al. 2019), suggesting alternative factors shaping bird beak evolution.

Here we investigate the relationships between beak integration, disparity, evolutionary rates and ecological specialization on the ‘megaevolutionary’ radiation of birds, given the importance of this biological innovation for the second largest extant tetrapod clade. We leverage vast resources for multidimensional trait information in the Mark My Bird (MMB) 3D beak landmark dataset (Cooney et al. 2017) and AVONET database (Tobias et al. 2022). Since its original description by Cooney et al. (2017), few studies have used the entire MMB sample for macroevolutionary analyses (but see Guillerme et al. 2023), likely due to computational costs of applying phylogenetic comparative methods to such massive datasets (>8000 species). To overcome these limitations, we present a novel methodological workflow for partitioning higher phylogenetic clades into order- or (passerine) family-level analyses, to understand how intrinsic factors shape beak macroevolutionary patterns. Specifically, we attempt to identify a causal network among these four major macroevolutionary ‘currencies’, to infer the potential impacts of ecological specialization on morphological integration, and their coordinated effects on rates of phenotypic evolution and disparity.

## RESULTS

### (a) Avian beak morphospace

Nearly all of modern avian beak diversity is captured in two dimensions (PC1-PC2 84.5%; Fig. 1C), with the majority of shape variation falling along a spectrum of beak depth-to-length ratios from hummingbirds (Trochilidae; negative PC1) to casque-headed species (positive PC1). PC2 describes variation in beak width and position of the nasofrontal hinge relative to gape (i.e., posterior tomial landmarks), ranging from birds with short, wide beaks and large gapes like nightjars (Caprimulgidae) and potoos (Nyctibiidae) on the negative side to those with thin beaks and casques or frontal shields at the positive extreme, such as Ross’s turaco (*Tauraco rossae*) and great hornbill (*Buceros bicornis*).

**Figure 1.**
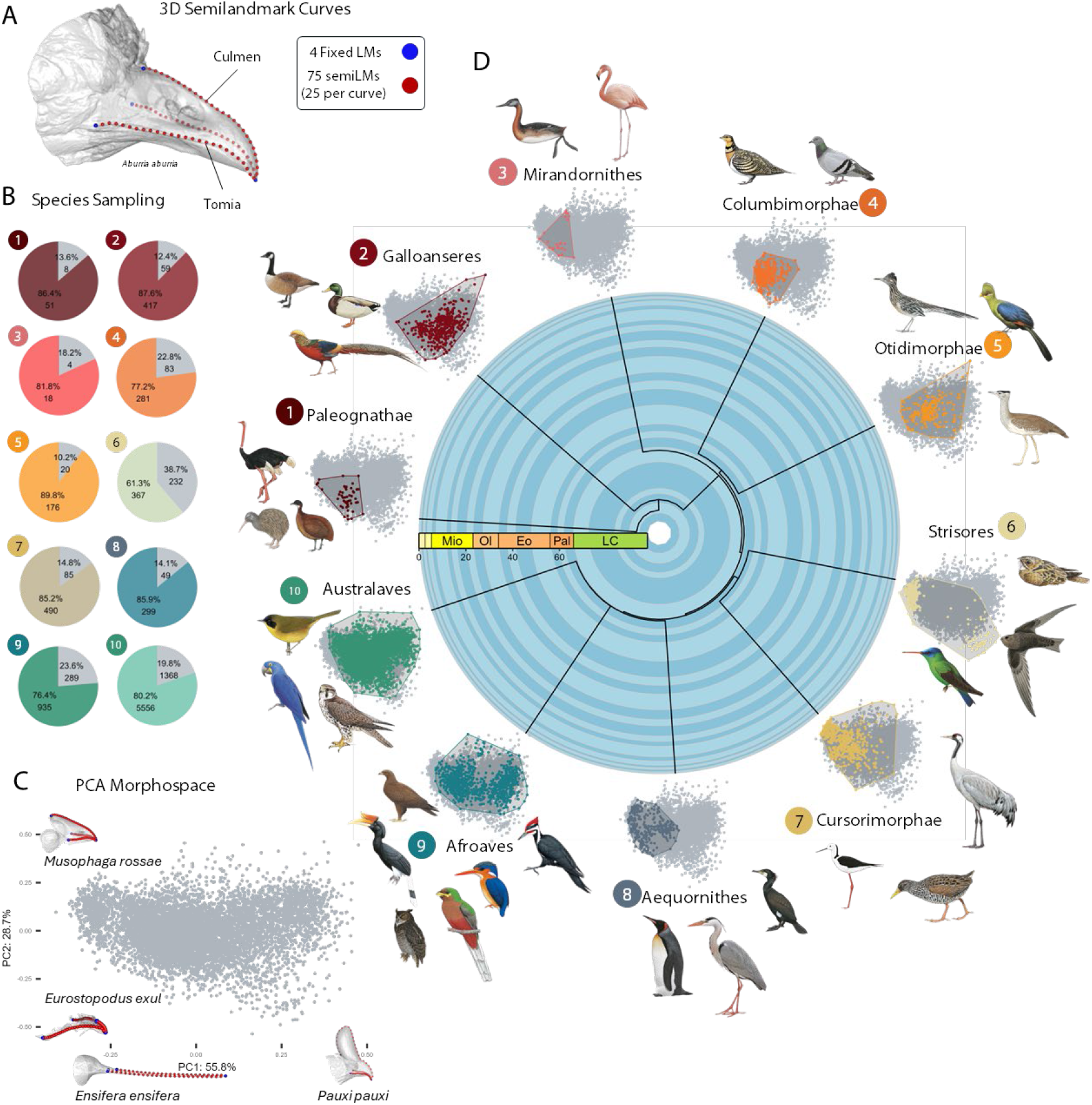
Summary of Mark My Bird (MMB) 3D landmark data on avian beaks. A) Four fixed/discrete landmarks (LMs) are shown in blue, connected by 25 semilandmarks along each of the three curves (red). B) Sampling per major clade according to Stiller et al. (2024), comparing the number of landmarked species to observed species richness, following Clements 2021 taxonomy (Clements et al. 2021). C) Bird beak morphospace along the first two principal components, containing over 84% of total shape variation. Examples of species from negative to positive shape extremes are shown for each axis. D) Morphospace occupation per major sampled clade, represented by color-coded convex hulls overlayed on the PC space of the entire MMB dataset, shown in gray. The dated phylogeny is simplified from McTavish et al. (2025). Bird illustrations courtesy of Cornell Lab of Ornithology | Birds of the World (https://doi.org/10.2173/bow).

Morphospace occupation among higher taxonomic groups (Fig. 1D) is widespread and largely overlapping, with the exception of Paleognathae, containing the largest and heaviest flightless birds (ostriches, emus, but also chicken-sized kiwis), and the neognath clades Mirandornithes (flamingos, grebes) and Aequornithes (water birds like penguins, pelicans and loons). Although the former two constitute the smallest sampled clades (59 and 22 species, respectively), over 81% of their diversity is represented in morphospace (Fig. 1B). Similarly, over 85% of Aequornithes is included in the MMB dataset (Fig. 1B), making sampling bias an unlikely explanation for the observed patterns. Rather, these clades contain birds with unique beak morphologies, typically falling along the negative edges of PC1-PC2 morphospace (Fig. 1D). For example, paleognaths lack the lateral bar present in ancestral theropods and neognaths (a bony load-bearing element connecting the posterior ends of the medial dorsal bar and laterally symmetrical ventral bars, captured by the culmen and tomial semilandmark curves, respectively; Fig. 1A), giving rise to elongate flat bills less adapted for complex tasks (Gussekloo et al. 2017). Within Mirandornithes, grebes (Podicipediformes) typically have long, pointed, laterally compressed beaks suited to their mainly aquatic diving lifestyles, although some species display short thick beaks adapted for crushing hard-shelled prey (e.g., pied-billed grebe, *Podilymbus podiceps*). Their sister taxon flamingos (Phoenicopteriformes) are characterized by downward-curved, L-shaped beaks with prominent lamellae along the edges, adapted for filter feeding in shallow water. In contrast, Aequornithes are a diverse clade of water-associated birds ranging from long-legged waders to diving specialists, with long, slender, pointed beaks for fishing (e.g., herons and egrets, Ardeidae), broad scoop-shaped beaks for filter feeding (pelicans, Pelicanidae), and robust hooked beaks for grabbing and tearing prey (cormorants, Phalacrocoracidae), hence their wider range along PC1.

### (b) Impacts of evolutionary allometry on beak shapes

Observed differences in morphospace occupation may be driven by allometry, for example if beak shape diversifies more with larger body size (e.g., ostriches, pelicans, flamingoes). However only 23 of the 90 subclades showed significant but negligeable allometry, with regression of beak shape on (log) centroid size resulting in mean Z-scores of 2.44 and mean R^2^ of 0.09 (see Table S5 (OLS) or Table S7 (PGLS) for full subclade results). Estimating Pagel’s λ per subclade from the residual landmarks (Table S6) returned a wide range of phylogenetic signal estimates, with the majority below 0.5 (mean λ = 0.26). After applying PGLS for allometric regressions, the number of subclades with significant allometry increased to 29, although still with relatively weak size-shape correlations (mean Z-score = 2.31, mean R^2^ = 0.12; Table S7). For all subclade phylogenetic allometric corrections, PGLS was the preferred model with the exceptions of Artamidae and Grallaridae, for which OLS was the better fit (see AICc scores, Table S8).

### (c) Variation of integration in orders and passerine families

Considerable variation in the strength of beak integration was recovered across subclades (Fig. 2A). Passeriformes covered the entire range of scaled effect sizes (ZR=0.46-3.15; Fig. 2B), with the lowest overall values belonging to Rhipiduridae (fantails) and the highest to Muscicapidae (Old World flycatchers). Both of these groups are insectivorous, with fantails having more flat and triangular beaks adapted to catching insects in flight. Columbimorphae (Pterocliformes and Columbiformes) also displayed a large range of ZR scores (0.65-2.7; Fig. 2A), despite its members having similar short, cone-shaped beaks (Özkan et al. 2024). Landmark integration was highest along the dorsal edge of the beak (Fig. 2 insets), where the nasofrontal landmark and posterior portion of the semilandmark curve showed the highest covariance. Those together with the posteriormost tomial landmarks also showed the largest standard deviations of covariances with all other landmarks, meaning that the strength of covariation varied across species within the same subclade – a pattern consistent in both passerine and non-passerine clades.

**Figure 2.**
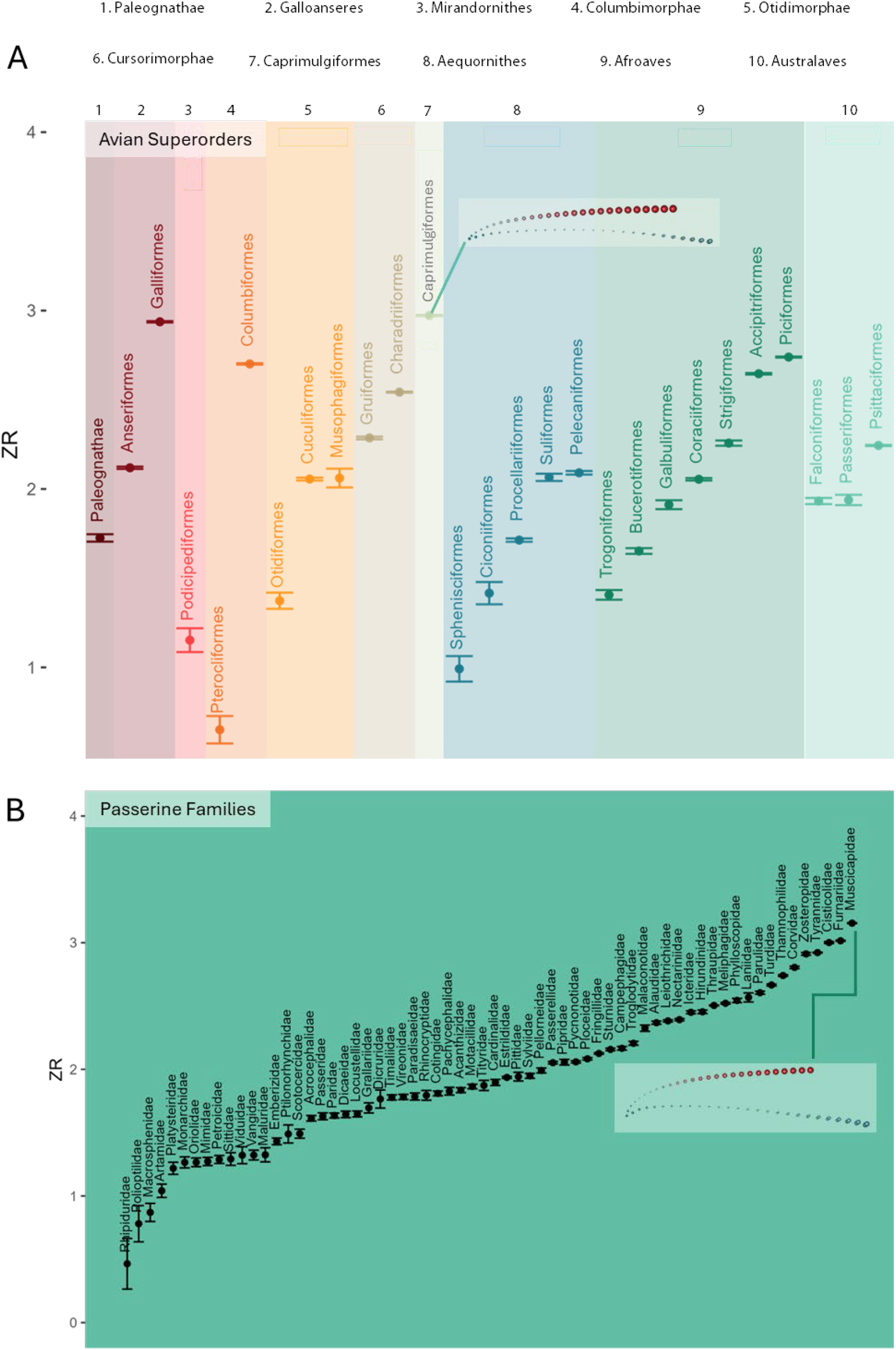
Degree of covariation among beak landmarks within and between A) major avian subclades and B) passerine families, calculated as effect size (Z score) of the observed relative eigenvalue index. Here, effect size is translated to a positive scale (ZR), such that ZR ranges from zero (no trait covariation) to positive values, indicating increasing levels of trait covariation (Conway and Adams 2022). Higher level clade colours match those in Fig. 1. Insets show examples of landmark integration on the consensus shape for the highest ZR scores in A) non-passerines (Caprimulgiformes) and B) passerines (Muscicapidae), with landmark colour indicating mean covariance with all other landmarks (redder = higher covariance, blue = lower), and landmark size indicating standard deviation of landmark covariance with all other landmarks.

### (d) Beak evolutionary rates and disparity

Beak disparity, measured as Procrustes variance, was positively and significantly correlated with BM rates, σ^2^ (R^2^ = 0.076, Z = 1.93, p = 0.019; Fig. S3A). Higher disparity was observed in non-Passerine orders than in Passerine families (Fig. S3B). The greatest disparity was found in Caprimulgiformes (nightjars, members of Strisores), which tend to occupy the negative quadrant of PC1-PC2 morphospace, while the lowest disparity was in Ciconiiformes (storks) and Aequornithes, in the upper left quadrant (Fig. 1D). Phylogenetic ANOVA between passerines and non-passerine orders suggest a significant difference in disparity (p = 4.091*e^-7^), even after accounting for clade age (CA) or species richness (SR) biases by using residuals from CA/SR PGLS and excluding CA/SR outliers (such as Caprimulgiformes). Rates, however, were relatively low across the subclade-level phylogeny, with a few lineages displaying independent bursts of elevated tempo, such as Galliformes, Falconiiformes, Cotingidae, Dicaeidae, Viduidae, Fringillidae, and Thraupidae (Fig. S3C). Phylogenetic ANOVA between passerines and non-passerine orders suggest no significant difference in rates (p = 0.8901; performed on residuals from clade age/species richness PGLS and excluding clade age (CA) and species richness (SR) outliers). Nonetheless, some passerine lineages experienced relatively elevated rates, where beaks of Phylloscopidae (leaf warblers) evolved an order of magnitude faster than the next fastest passerine family Scotocercidae (bush warblers) (Table S11).

### e) Ecological specialization estimated by the Gini coefficient (GC)

Birds display the full range of trophic niche specialization, with passerines distributed across the entire spectrum (Fig. 3). Estimates of GC with and without phylogenetic history (i.e., simulated proportions of time spent in trophic states via stochastic character mapping (SCM)) resulted in highly similar values (Fig. 3A), however a t-test and Wilcoxan signed-rank test revealed significant differences (p = 0.0129; p < 0.0001). Subclades that differed most among the two methods were the passerine families Mimidae, Emberizidae, Oriolidae, Ptilorhynchidae and Pachycephalidae, and the order Pelececaniformes (Fig. 3B). The simple GC showed higher phylogenetic signal (K = 0.32) than SCM-based (K = 0.24), and was marginally positively correlated with species richness (R^2^ = 0.04, p = 0.059) while SCM-based GC was not; neither was correlated with clade age (Table S9). According to SCM simulations, the lowest GCs belong to Oriolidae (Old World orioles, GC = 0.015), Emberizidae (buntings, GC = 0.017) and Mimidae (New World thrashers, mockingbirds and relatives, GC = 0.019), with near-equal times spent across trophic niches. Many subclades had GC = 1, indicating that all members occupied a single dietary category throughout their evolutionary history. Of those subclades whose GC did not equal one, Furnariidae (ovenbirds and woodcreepers, mainly insectivores with few aquatic predators) had the highest degree of specialization (GC = 0.997; Table S11).

**Figure 3.**
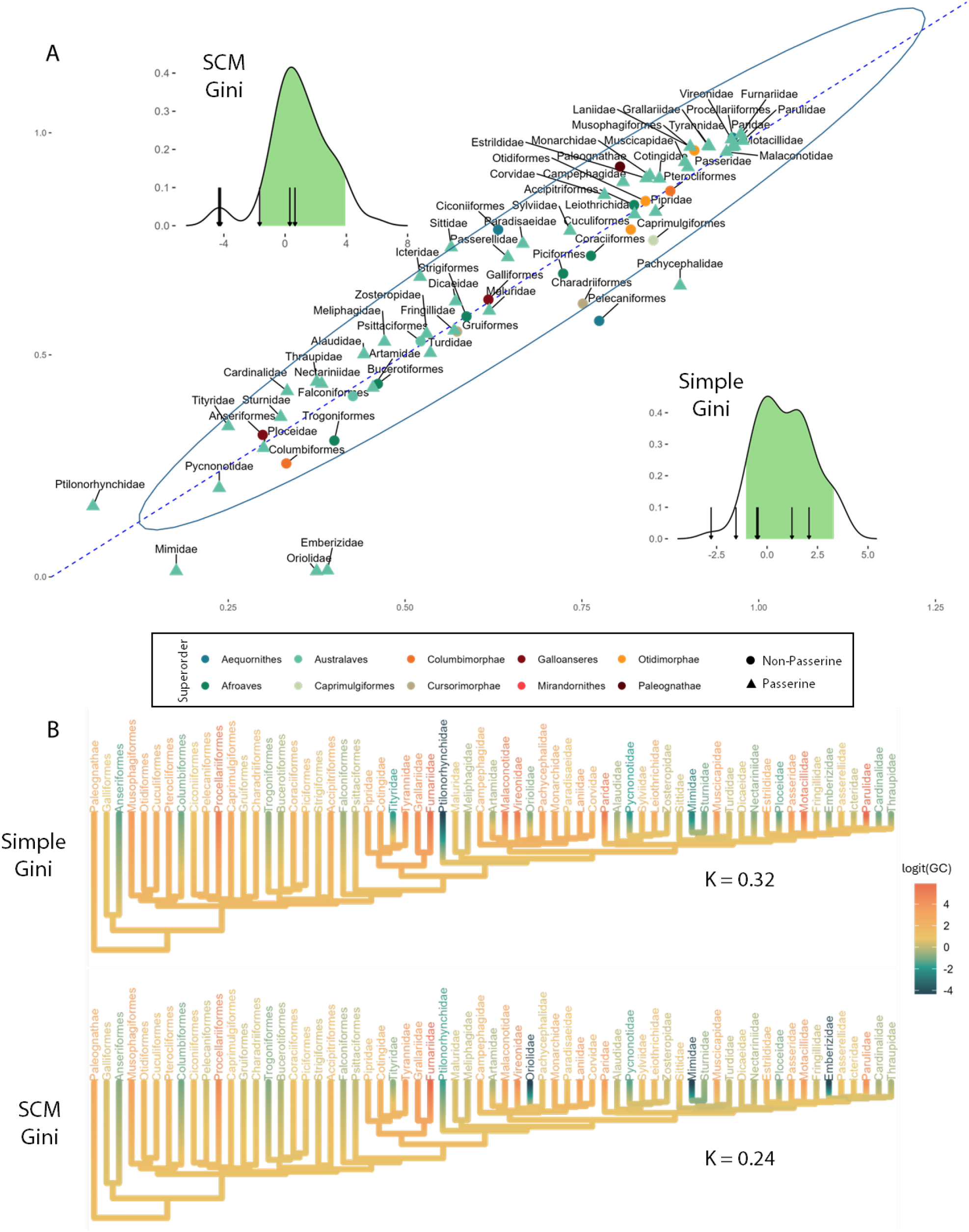
Properties of simple versus stochastic character mapping (SCM)-based Gini coefficients (GC). A) Scatterplot of the two estimates for each subclade, with 95% confidence interval ellipsoid. Density plots shows distributions of simple and SCM GC estimates per subclade. Vertical arrows point to outlier subclades falling outside of the ellipsoid, denoted in green. B) Continuous character mapping of simple CG vs SCM-based GC, based on Brownian motion model assumptions. Blomberg’s K reported for evaluation of phylogenetic signal for both. Hotter colors represent higher logit-transformed GC value, meaning relatively more specialized clades. Cooler colors represent lower logit-transformed GC values, indicating more generalist clades.

### (f) Relationships among the four macroevolutionary currencies

We first assessed the effects of clade age, species richness, and their interaction on the four major macroevolutionary variables (Fig. S2; Table S10).Correlations between integration and clade age or species richness were non-significant, with small effect sizes (Table S10). Variation in evolutionary rates was best described by clade age, with beaks of older lineages evolving significantly slower than younger ones (Fig. S2; Table S10). Disparity was positively and significantly associated with species richness (the preferred model according to AIC_c_ values; Table S11) but also clade age and their interaction, with the most disparate beak shapes belonging to the largest sampled subclade Caprimulgiformes (303 species). However, note that the next-most disparate subclade Sylviidae (warblers) comprised only 55 sampled species – roughly average for our dataset (Table S3). The best fitting descriptor for ecological specialization was also species richness, with smaller clades spending unequal proportions of time across fewer trophic niches, i.e., being more specialized during their evolutionary history; clade age had no effect (Fig. S2; Table S10).

Outlier clades removed based on clade age and species richness from downstream analyses were: Paleognathae, Anseriformes, Musophagiformes, Caprimulgiformes (Strisores), Accipitriformes, Phylloscopidae, Polioptilidae, Pterocliformes, Psittaciformes, Dicruridae, Oriolidae, Mimidae, Emberizidae, Scotocercidae, Ciconiiformes, Sylviidae and Paradisaeidae. Among the remaining 73 subclades, we found no effect of ecological specialization on integration (Table 1; Fig. 4A). In the best fitting model for rates of phenotypic evolution (represented by the multivariate BM rate σ^2^), both integration and ecological specialization were significant predictors (Fig. 4A,B), although their interaction had no effect (Table 1). Clades evolving at faster rates had stronger beak integration and were more generalist in their dietary habits (Fig. S11), meaning equal proportions of times spent in their past among different feeding strategies. Neither integration nor its interaction with ecological specialization were significantly associated with beak disparity, although specialization on its own had a relatively large effect size (Table 1; Fig. 4D), suggesting that generalist lineages tend to have more diverse (disparate) beak shapes. See Figs. S4-S7 for scatterplots between non-residual macroevolutionary variables discussed above, and Figs. S8-S10 for scatterplots of residual values, after accounting for clade age and species richness.

**Table 1.** PGLS results between the four macroevolutionary currencies. Change in AIC value (ΔAIC) was calculated for each model except for Integration ∼ Ecological specialization, as no competing models were tested. Preferred models are shown in bold, with the summary statistics for each predictor variable given for best fitting rates model.

| PGLS MODEL | R<sup>2</sup> | P | F | Z | AIC<sub>c</sub> | ΔAIC<sub>c</sub> |
| --- | --- | --- | --- | --- | --- | --- |
| Integration ~ Ecological Specialization | 0.012 | 0.359 | 0.840 | 0.414 | — | — |
| ***Rates ~ Integration + Ecological Specialization + Integration\*Ecological Specialization*** | **0.185** | **0.001** | **6.201** | **2.963** | **115.753** | **0** |
| *Integration* | *0.078* | *0.01* | *6.82* | *2.09* | — | — |
| *Ecological Specialization* | *0.13* | *0.002* | *11.72* | *2.67* | — | — |
| *Integration\*Ecological Specialization* | *0.0007* | *0.81* | *0.061* | *-0.93* | — | — |
| Rates ~ Integration | 0.078 | 0.016 | 5.995 | 1.951 | 123.262 | 7.508 |
| Rates ~ Ecological Specialization | 0.111 | 0.005 | 8.883 | 2.328 | 120.574 | 4.821 |
| Disparity ~ Integration + Ecological<br>Specialization + Integration*Ecological<br>Specialization | 0.015 | 0.329 | 1.166 | 3.834 | 77.565 | 3.782 |
| Disparity ~ Integration | 0.002 | 0.706 | 0.149 | 4.385 | 77.006 | 3.222 |
| **Disparity ~ Ecological Specialization** | **0.045** | **0.070** | **3.364** | **5.103** | **73.784** | **0** |

**Figure 4.**
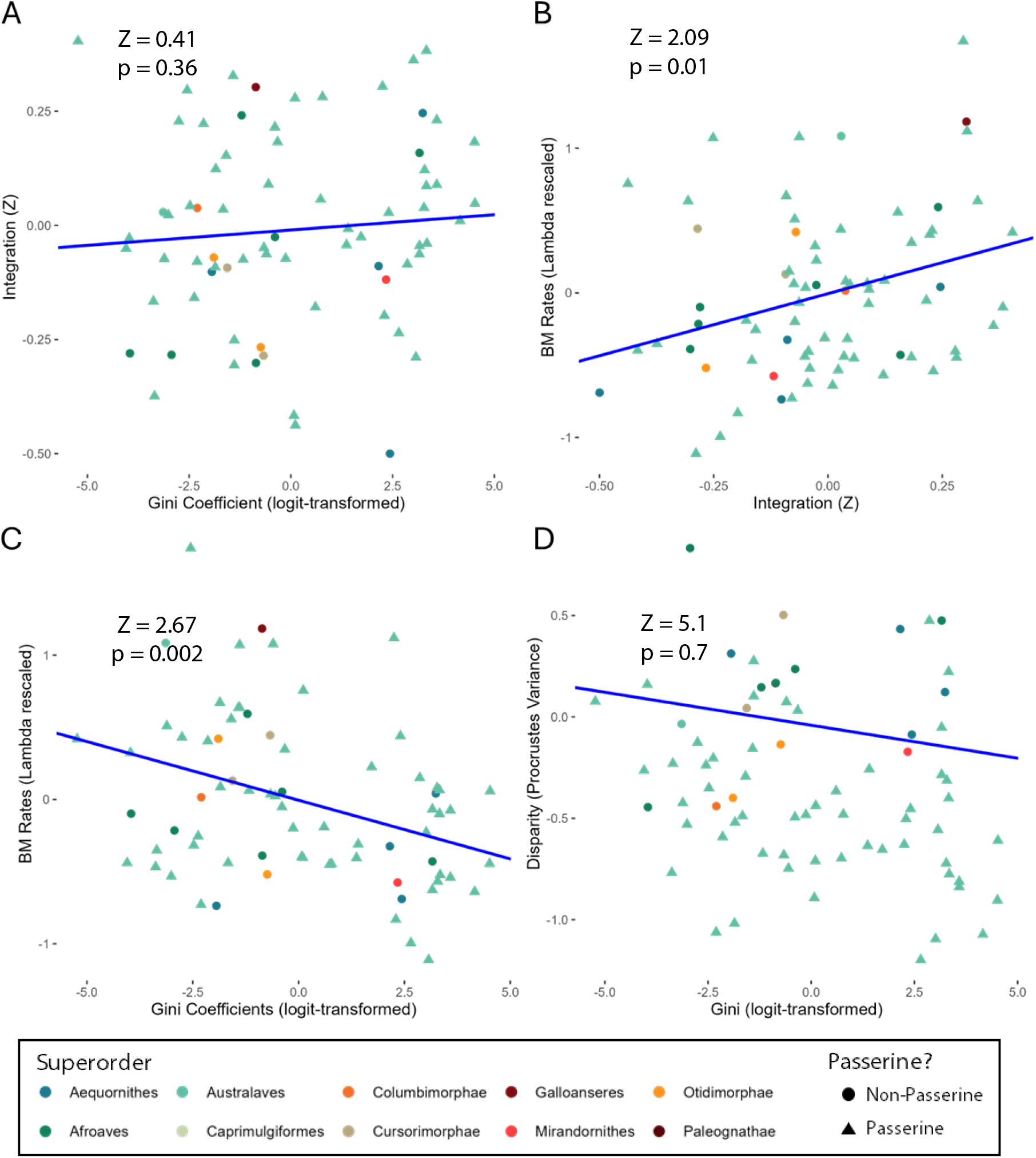
Main PGLS results of A) Integration on ecological specialization. B) Rates on integration. C) Rates on ecological specialization. D) Disparity on ecological specialization. Points are color-coded according to major clades of Stiller et al. (2024) (see Fig. 1).

## DISCUSSION

The transformation of birds from theropod dinosaurs is marked by reconfiguration of the bony snout into a lightweight kinetic and edentulous beak, enabling diverse biological functions (Lautenschlager et al. 2013; Bhullar et al. 2016; Wang et al. 2017b; Garland et al. 2025). Together with flight, this phenotypic innovation promoted adaptive radiations into numerous ecological niches (Cooney et al. 2017; Chira et al. 2020), giving rise to an astounding array of beak morphologies linked to specific feeding strategies, from the flower-probing tubular bills of hummingbirds, stout seed-crushing bills of finches, broad fish-scooping beaks of pelicans to the hooked flesh-tearing beaks of raptors. This remarkable diversity implies a lack of constraints on the developmental processes underlying beak formation, potentially reflected in the strength of relationships among multivariate beak traits.

We quantified variation among 79 three-dimensional landmarks on the upper beak surface of 8627 species, showing that the strength of integration, or landmark covariance, varies widely across avian subclades and is independent of beak size, phylogeny, clade age and species richness. Lineages with faster rates of phenotypic evolution have more strongly integrated beaks and more diverse trophic histories, indicating that beak integration facilitates avian evolution. This pattern contrasts with predictions that higher trait covariance hinders phenotypic change, by preventing correlated traits from responding to differential selection pressures (Monteiro and Nogueira 2010; Hedrick et al. 2020; Sansalone et al. 2022). Instead, we find that integration among upper beak regions likely promotes rapid phenotypic diversification along the major axes of variation (i.e., PC1-PC2; Fig. 1C), favoring trait combinations that maximize performance across multiple trophic niches. These so-called ‘lines of least resistance’ (Schluter 1996; Guillerme et al. 2023) emphasize the role of genetic and developmental processes in directing beak divergence, which across the megaevolutionary radiation of birds appears independent of allometry – an otherwise strong driver of morphological integration (Klingenberg 2016).

These intrinsic factors do not interact directly, but likely amplify each other under opportunistic conditions. For example, lineages spending more of their evolutionary history among different trophic niches (i.e., lower GC) evolved faster when their beaks were more integrated (Fig. 4; Fig. S11), and tended to reach higher morphological disparity (Fig. 4). However ecological specialization itself did not predict integration, nor did integration produce more disparate beak shapes (Table 1). Together this pattern suggests that beaks diversify into multifunctional tools by maintaining coordination among key traits, which in turns leads to rapid evolution as lineages expand into unoccupied niches. This may explain why we recover lower rates among more specialized clades (e.g., Furnariidae, Thamnophilidae;, Fig. 4C; Figs. S9B, S10B); stabilizing selection around adaptive peaks may limit the tempo of evolution once lineages fix on that optimum (Hansen 1997). However, as environments change and new niches open, bouts of rapid evolution can occur during niche shifts, whereby generalists, which experience weaker stabilizing selection and therefore maintain greater standing phenotypic or genetic variation, respond faster to ecological opportunity (Dennis et al. 2011; Cooney et al. 2017; Hayes et al. 2024).

This scenario appears to have played out early in the history of neornithine birds, which famously underwent early burst dynamics of trait evolution following the KPg extinction (Harmon et al. 2010, Feduccia 2014). Thus, older lineages experienced adequate time to converge on adaptive peaks, after which diversification decelerated as species crowded into increasingly saturated niche space (Cooney et al. 2017). We recovered slower rates among older clades which were also more phenotypically diverse (Fig. S2), further supporting this hypothesis. Interestingly, the basal Palaeognathae and Mirandornithes were notably depauperate in morphospace occupation (Fig. 1D). While members of these clades have usual bill shapes (see Results section a: Avian beak morphospace), they also underwent significant extinctions since their origins in the late Cretaceous, with the fossil record documenting beak/feeding morphologies no longer present in the modern forms (Allentoft et al. 2014; Agnolin 2017; Hansford and Turvey 2018; du Toit et al. 2020; Mayr 2022; Widrig and Field 2022). Therefore integration does not guarantee evolutionary stability, but may produce novel phenotypes and ecologies that become outcompeted or obsolete as environments change. This is evident for many avian lineages, as end-Cretaceous events may have driven extinction of toothed and specialized Mesozoic birds (Larson et al. 2016), palaeognath megafauna (moas, elephant birds) were largely lost after human colonization (Allentoft et al. 2014; Hansford and Turvey 2018), and many extinct seabirds and stem flamingo and grebe relatives declined with Cenozoic oceanic and habitat shifts (Mayr 2004; Ksepka et al. 2013; Torres et al. 2015) (see Supplementary Section II for further discussion of morphospace patterns for recently phylogenomically-resolved clades).

The strongest landmark covariance was found along the posterior culmen, while ventral and anterior beak regions were less tightly coordinated (Fig. 2 insets). Although the upper bird beak is typically considered a single skeletal unit, the initial beak primordia develops from two distinct tissue condensations: the prenasal cartilage (chrondrocranium), followed by the premaxillary bone (dermatocranium) (Mallarino et al. 2011) . These are regulated by different sets of molecules, whose heterochronic expression during embryonic development can alter beak growth along independent (depth/width vs length) dimensions (Abzhanov et al. 2004, 2006). Although we did not consider developmental modules in our analysis, the region of highest landmark covariance concentrates on the premaxilla where it contacts the skull roof and adjacent bones (insets, Fig 2). This junction is critical for transferring forces from the bill tip to the skull during feeding (Bock 1999; Soons et al. 2010; Cuff et al. 2015; Gussekloo et al. 2017; Plateau and Foth 2020), enabling cranial kinesis (Stefanini et al. 2025), and is a key growth zone boundary during craniofacial morphogenesis (Marcucio et al. 2005; Hu and Marcucio 2009).Experimental studies demonstrate that higher levels of regulatory genes expressed in this region alter premaxillary elongation, depth and dorsal profile, thus driving the extensive beak variation observed among Darwin’s finches (Abzhanov et al. 2004, 2006; Mallarino et al. 2011, 2012). We therefore aim to evaluate beak morphology in light of developmental modularity in the future, by sampling more complete 3D shapes (e.g., using surface pseudolandmarks) as well as internal structures/sutures between bones. Ideally these data can be combined with gene expression and functional analyses (Mallarino et al. 2011, 2012; Martínez-Abadías et al. 2016; Cheng et al. 2020; Huang et al. 2022), to better understand the factors driving beak differences within and among species.

We focused on a single ecological variable measured by the Gini coefficient (trophic niche), however other traits in the AVONET database (Tobias et al. 2022) warrant exploration in relation to beak macroevolutionary patterns. The bird beak is highly multifunctional, such that factors including habitat type, lifestyle (e.g., primarily aerial, perching, aquatic, terrestrial), and migratory behavior likely impose concurrent selection pressures on morphology. Indeed, several nondietary traits have been shown to play important roles in determining beak shape in specific groups, for example size and integration with the braincase in raptors (Bright et al. 2016), vocal signal structure (Podos 2001), and thermoregulation (Friedman et al. 2019). Nevertheless, we found that birds display the full range of dietary specialisation as simulated by stochastic character mapping, which differed significantly from extant patterns and was independent of species richness (Fig. 4A,D; Fig. S2). This may represent an advantage of SCM-based GC over the simple version, in that it is less biased by sample size and phylogenetic relatedness. Our method is readily applicable to other ecological traits across the tree of life, and to proportions of evolutionary time spent among different developmental states such as numbers of digits (Cooper et al. 2014) or tooth cusps (Burroughs 2019) to infer ancestral shifts in developmental patterning.

The lability of avian beak covariance was evaluated by Guillerme et al. (2023), who decomposed its evolutionary patterns into two modes: elaboration (evolution along a clade’s major axis of phenotypic variation) and innovation (evolution away from that axis). Over half (15/27) of avian orders appeared to innovate at the expense of elaboration, while passerine families displayed greater levels of elaboration (Guillerme et al. 2023). The majority of points in our analyses are passerines (64/90 subclades), potentially introducing higher correlations between rates and integration, assuming elaboration is a product of higher trait covariance (i.e., evolution along lines of least resistance) (Schluter 1996; Marroig and Cheverud 2005; Guillerme et al. 2023). Adaptive radiation theory predicts that substantial time is needed for a clade to realign its primary axes of evolutionary covariance, suggesting that Passeriformes, being on average younger than other clades (Table S2), may not have accumulated sufficient time or ecological opportunity to reduce developmental biases or shift away from their adaptive optima.

## CONCLUSIONS

We found that stronger integration of the beak is associated with faster rates of phenotypic evolution across avian subclades, also associated with low ecological specialisation. Although theoretical expectations suggest higher integration should constrain evolution (Cheverud 1982), our results are consistent with the ‘facilitation hypothesis’ (Goswami and Polly 2010a), whereby certain macroevolutionary variables are enhanced by functional trait covariance along the lines of least resistance. Most studies have evaluated the relationship between integration and disparity, but not rates (Goswami and Polly 2010a). Here we broaden expectations of the facilitation hypothesis to include rates of evolution, being proportional to the magnitude of integration. This counters previous findings that integration does not influence rates of evolution in carnivoran (Goswami et al. 2014) and avian crania (Felice et al. 2018). While those studies estimated rates and integration per anatomical module, we estimated macroevolutionary variables per subclade within the upper bird beak (i.e., a single module). The striking variation in integration values recovered across bird orders and passerine families highlights the complex developmental history of this deceptively simple element, derived from multiple cranial bones, tissue types and molecular pathways. We also provide the first detailed description of avian beak morphospace using the full MMB dataset, applying the recently resolved family-level genomic phylogeny to explore macroevolutionary patterns. Future analyses should harness this integrated resource together with developmental models and gene expression data to further explore the underlying factors shaping bird beak morphogenesis, to build an optimized model for the growth and diversification of one of nature’s most efficient tools.

## MATERIALS AND METHODS

### (a) Phylogenetic comparative dataset

We described beak shape using 3D landmark data from the MarkMyBird.org citizen science project (Fig. 1A; Cooney et al. 2017), comprising 8748 species under the BirdTree taxonomic classification (Jetz et al. 2012) or 8627 species under the eBird taxonomic classification (Sullivan et al. 2009), thus representing ∼ 80% of modern avian diversity (Fig. 1B). The landmark scheme (Fig. 1A) is composed of four anatomical landmarks placed on 3D surface meshes of each bird beak, one at the anteriormost point of the beak, one at the posteriormost point coinciding with the nasofrontal joint, and two along the posteriormost points of the lateral beak (tomia). These fixed landmarks demarcate two semilandmark curves along the tomial edges and one along the culmen (dorsal edge of the beak), with 25 semilandmarks per curve. Landmarks were subjected to Procrustes superimposition to standardize for scaling, orientation, and rotation among specimens using the *gpagen* function in geomorph 4.0.1 (Baken et al. 2021), specifying semilandmarks along each curve as sliding. We performed principal components analysis (PCA) of the landmark data to visualize variation in beak morphospace, and to compare clade-specific patterns of morphospace occupation that may relate to ecological specialization.

Disparity, rates, integration, and ecological specialization were analyzed per taxonomic subgroup, using a recent supertree of 10,824 bird species (McTavish et al. 2025). This supertree corresponds to the Clements 2021 taxonomic system, with a backbone topology heavily weighed in favor of the recent family-level genome phylogeny of Stiller et al. (2024). The latter study presented the first highly resolved and well-supported clade relationships for previously contentious phylogenetic placements of major bird subclades, such as Mirandornithes, Columbaves (Columbimorphae, Otidimorphae), Telluraves, and the novel clade Elementaves (Aequornithes, Phaethontimorphae, Strisores, Cursorimorphae, Opisthocomiformes). We extracted the supertree through the R package clootl v0.1.1 using the function *extractTree* (all species, Clements version 2021) (Miller et al. 2025). See Supplementary Section Ia for full details on standardizing taxonomic systems between datasets.

All ecological and evolutionary estimates were performed by collapsing the phylogeny into subclades with 15 or more species defined by the major non-passerine order-level (Table S1) and passerine family-level divisions (Table S2) in McTavish et al. (2025). All five orders in Palaeognathae, however, were grouped as a single subclade due to low species richness of some of its orders (e.g., Struthioniformes, 2 species). Relabelling paleognath orders into one clade dropped the number of orders analyzed from 40 to 35 (excluding Passeriformes), resulting in a total of 90 subclades. See Supplementary Section Ib for details on producing the 90-tip subclade-level phylogeny, and Tables S1-S4 for subclade summary data, including clade age and species richness.

### (b) Correction for allometric and phylogenetic signal

Shape covariation with size has demonstrably affected inferences of various macroevolutionary processes, including morphological integration (Klingenberg 2016). Since size is thought to impact the whole organism, allometry introduces a global correlative factor, inflating overall shape covariance and potentially masking more subtle covariance patterns. Significant impacts of allometry in bird beaks have revealed weak influences in previous analyses (Rombaut et al. 2022). Nonetheless, we ran Procrustes regression with whole landmarks as the response variable and centroid size as the predictor for each subclade. We extracted residual landmark estimates corrected for allometry and phylogenetic signal following the protocol of Natale & Slater (2022). This approach expects the residual error structure of the landmark data to be described by Pagel’s λ assumptions as opposed to Brownian motion (BM) (Natale and Slater 2022). To estimate λ from residual landmarks from an ordinary least squares (OLS) Procrustes regression of landmarks onto (log) centroid size (see Table S5 for OLS summary statistics), we used the penalized likelihood procedure via the function *fit_t_pl* in RPANDA v2.5 (Morlon et al. 2016) for high dimensional datasets, extracting the best-fitting λ per subclade (Table S6) estimate for calculation of a phylogenetic covariance matrix used directly in a Procrustes regression in *procD*.*lm* in geomorph (provided under argument *Cov*; Adams and Collyer 2018; Baken et al. 2021). See Table S7 for summary statistics from PGLS with λ-rescaled covariance matrices. Using *procD*.*lm* with a custom phylogenetic covariance matrix in this way is equivalent to phylogenetic generalized least squares (PGLS), but allows relaxation of the default assumption of BM that *procD*.*pgls* uses to obtain the phylogenetic transformation matrix (see Natale and Slater 2022 for discussion). For each subclade, we chose residuals from either OLS or PGLS models on the basis of AIC_c_ (Table S8). All regression analyses were run for 10,000 permutations. See Supplementary Section Ic for all analytical details.

### (c) Bird beak integration

Several measures of integration are available for multivariate trait data (see Goswami and Polly 2010b for extensive review). Eigenvalue variance is arguably the most intuitive, as it is based on the dispersion of eigenvalues derived from a multivariate covariance matrix (Wagner 1984; Pavlicev et al. 2009). If covariance among traits is greater than within-trait correlation (or within subsets of traits, i.e., modules), eigenvalue variance will be greater. Conversely, eigenvalue variance will be lower if greater covariance occurs within individual traits or modules (Wagner 1984; Pavlicev et al. 2009). Therefore, greater eigenvalue variance indicates greater integration among traits. For each subclade, eigenvalue variance standardized to effect size (Z_Vrel_; Conaway and Adams 2022) was used for comparisons of the strength of beak landmark integration across bird subclades. Z_Vrel_ was calculated via the function *integration*.*Vrel* in geomorph (Baken et al. 2021), using pruned subclade-level phylogenetic trees (Pagel’s λ rescaled) as input to account for phylogenetic signal (Adams and Felice 2014).

### (d) Phenotypic rates and disparity

We estimated multivariate BM rates of evolution (σ^2^) per subclade using sigma.d (function modified from code available in Appendix 2 of Adams (2014)). These estimates are robust to Type 1 error rates and display high power compared to methods using standard evolutionary rate matrices, the latter of which can lead to unstable likelihood calculations when the number of traits exceeds the number of species (Adams 2014). This property is desirable when analyzing subclade level partitions of landmarks, where in most subclades the trait dimensions exceed the number of taxa (79 landmarks * 3 dimensions = 237 variables). Prior to rate estimation, we used the set of best fitting Pagel’s λ values per subclade to transform each subclade phylogeny.

Procrustes variance is often treated as an adequate measure of morphological disparity for landmark data, calculated as the trace of the covariance matrix (sum of variances) divided by the number of observations (Zelditch et al. 2012). Most works have utilized Procrustes variance among landmarks in the absence of correction for phylogenetic signal. As a solution, *morphol*.*disparity* in geomorph (Baken et al. 2021) allows direct input of PGLS models, where residual landmark estimates were extracted. We used PGLS outputs from our allometric and phylogenetic correction procedure to calculate Procrustes variance per subclade.

### (e) Ecological specialization

A variety of definitions make direct quantification of ecological specialization challenging (Devictor et al. 2010), however here we consider specialization in the framework of relative proportions of discrete ecological states. A simple measure of ecological specialization is to calculate the observed proportions of each ecological state across a sample of species (Levins 1968), usually done along (extant) tips of a phylogeny. However, such an approach does not account for the evolution of ecological state proportions through time. We therefore employ stochastic character mapping (SCM; Bollback 2006) to simulate alternative ecological state histories according to best-fitting discrete character evolutionary models (Mk models; Lewis 2001).

Following this approach, one can calculate the total amount of evolutionary time (time-calibrated phylogenetic branch length) spent in each discrete state, of which relative proportions indicate the degree of evolutionary ecological specialization. Several summary statistics of statistical dispersion exist (Devictor et al. 2010), of which the Gini coefficient (GC; Gini 1921), originally used to measure economic inequality, can be applied to ecological contexts (see Morelli et al. 2019). GC values range from 0-1, with zero indicating complete generalism where each ecological state receives equal proportions of evolutionary time spent in a subclade. If a subclade spends all its time in a single state, meaning it has specialized on that state during its entire evolutionary history, GC will equal one. Visually, GC may be calculated as the area between the Lorenz Curve, which represents the cumulative proportions of each state, and a one-to-one diagonal line representing perfect ecological generalism where each state has equal proportions of total time spent (Fig. S1).

In this work, we introduce a new method for estimating ecological specialization by applying GC to simulated times spent in categorical ecological states. Simulated times were produced via SCM on subclade phylogenies with discrete dietary characters. We focused on the ‘Trophic Niche’ categorical variable in the AVONET database (Tobias et al. 2022). The original Trophic Niche variable contains 10 dietary categories: vertivore, invertivore, scavenger, omnivore, aquatic predator, frugivore, aquatic herbivore, terrestrial herbivore, nectarivore and granivore. Species described as scavengers were placed into the vertivore category (species obtaining at least 60% of food resources from vertebrate animals in terrestrial systems, including mammals, birds, reptiles, etc.; Tobias et al. 2022), due to low sample sizes in our dataset (22 species).

Before applying SCM, we extracted subclade phylogenies from the avian supertree with corresponding AVONET trophic niche states for each species (Fig. S1: step 1). We then fit three models of discrete state evolution: Equal Rates (classic Mk model (Lewis 2001a)), Symmetric Rates, and All Rates Different (Paradis and Schliep 2019) to each subclade and its set of trophic niches (Fig. S1: step 2). We selected the best fitting discrete state transition model for each subclade via AIC_c_ scores for downstream SCM. For SCM simulations, we used the *fastSimmap* function in ratematrix v1.2.4 (Caetano and Harmon 2017) to produce 1000 character maps per subclade (Fig. S1: step 3). *fastSimmap* outputs times spent in each state per branch, from which the total amount of time the subclade spends in each state was calculated (Fig. S1: step 4). The GC was applied to the total times spent in each state via the function *Gini* in DescTools 0.99.6 R package (Signorell 2025), calculated for each SCM. The mean GC was then calculated from all stochastic mapping simulations per subclade. Prior to performing stochastic character simulations, we automatically assigned a GC equal to one if all species in a subclade had the same state. For most downstream analyses including PGLS (Grafen and Hamilton 1997), we used logit-transformed GCs to help normalize values naturally constrained to a 0-1 scale (Seiffert et al. 2024), using the *logit* function in the package car v3.1-3 (Fox and Weisberg 2018).

Finally, we compared our results to the simple (non-phylogenetic) GC estimates to detect potential advantages of the above approach. We also evaluated the effects of phylogenetic signal via Blomberg’s K (Blomberg et al. 2003), clade age, and species richness for each GC (Table S9, Fig. 3).

### (f) Linear regressions

We applied PGLS to subclade metrics for phylogenetically-corrected integration, phenotypic rates, disparity, and ecological specialization to test their evolutionary associations. The subclade level phylogeny of 90 tips (orders + passerine families) was used in all analyses, pruned from McTavish et al. (2025). We followed the protocol in Natale & Slater (2022), first using OLS residuals subjected to model fitting via Pagel’s λ. The λ-rescaled phylogeny was converted to a phylogenetic covariance matrix used in the PGLS residual error term to account for phylogenetic signal. For either OLS or PGLS, we used *procD*.*lm* in geomorph 4.0.1 with the covariance matrix provided under argument *Cov* (Baken et al. 2021), run for 10,000 permutations.

For many macroevolutionary variables, species richness and subclade age are important covariates which could bias PGLS inferences. We therefore tested the impacts of clade age and species richness on rates, disparity, integration, and ecological specialization individually. PGLS was performed using *procD*.*lm* on each of the four variables with either clade age, species richness, or an interaction between clade age and species richness, specifying a phylogenetic covariance matrix from the best fitting Pagel’s λ between the predictor and response variables. On the massive macroevolutionary scale represented by birds, a handful of lineages may experience differential macroevolutionary dynamics than the norm for the entire clade (Slater et al. 2010; Slater and Pennell 2014). We removed these ‘outliers’ from downstream analyses by detecting which subclades lie outside the 95% confidence interval ellipses for each PGLS regression using the *pointstoEllipsoid* function the package SIBER v2.1.9 (Jackson 2023). Scatterplots of all 90 subclades with their respective 95% confidence interval ellipses are presented in Figs. S3-S6.

PGLS analyses were performed between integration as a response variable and ecological specialization as predictor variable. We then assessed the role of integration, ecological specialization, and their interaction as predictor variables on disparity and rates. We report standard summary statistics (p-value, F, Z, AIC_c_, R^2^), assessing the best-fitting model with the lowest AIC_c_ value. For all analyses, we used the function *procD*.*lm* in geomorph 4.0.1 (Baken et al. 2021) for OLS and PGLS implementation with a phylogenetic covariance matrix estimated from rescaled tree branches according to Pagel’s λ, used to explain the residual error matrix (Pagel 1997). For the three term model, SS type was specified as Type II (Adams and Collyer 2018). Scatterplots of the best fitting variable pairs based on PGLS results are shown in Figs. S7-S9.

## Acknowledgments

We are grateful to Gavin Thomas (University of Sheffield) for access to the Mark My Bird landmarks and meshes; Center for Computational Evolutionary Morphometry (CCEM) at the University of Copenhagen; Rasmus Nielsen (CCEM) for guidance and funding; Michael Severinsen (CCEM) for ; Mike Collyer (Chatham University) and Dean Adams (Iowa State University) for training through their workshop “Geometric Morphometrics for R”.

## Funding

this work was supported by the Villum Fonden grant 00040582 (S.H.S).

## Author contributions

Conceptualization: R.C.E., C.A.H

Methodology: R.C.E., C.A.H.

Investigation: R.C.E.

Visualization: R.C.E., C.A.H.

Supervision: C.A.H., S.H.S.

Writing—original draft: R.C.E., C.A.H.

Writing—review & editing: R.C.E., C.A.H., S.H.S.

## Competing interests

all authors declare no competing interests.

## Data, code, and materials availability

please contact Gavin Thomas (University of Sheffield) to obtain MMB landmark data. Supplementary code and data necessary to reproduce all analyses are available in the following Zenodo repository: 10.5281/zenodo.19608071.

## Notes

https://doi.org/10.5281/zenodo.19608071

### Competing Interest Statement

The authors have declared no competing interest.

