## Supplementary Materials for "Morphological integration of the avian beak facilitates evolution along lines of least resistance"

Ely *et al.*

**This PDF file includes:**

Supplementary Text
Figs. S1 to S12
Tables S1 to S11

**Supplementary Text**

Taxonomic System Standardization

All species names in the phylogeny (McTavish et al. 2025), Mark My Bird (MMB) landmark dataset (Cooney et al. 2017), and ecological data represented in the AVONET database (Tobias et al. 2022) were standardized to match the phylogeny of McTavish et al. (2025), according to the Clements 2021 taxonomic system (Clements et al. 2021). The chosen supertree via McTavish et al. (2025) follows this system, such that only the MMB and AVONET datasets were modified. This resulted in an apparent drop in species numbers of the MMB dataset, from 8748 to 8627 species, when in fact these were equivalent due to the presence of taxonomic synonyms. To account for the presence of more synonymous species in the MMB dataset than in the Clements 2021 system, landmarks from the MMB dataset pertaining to equivalent species in the Clements 2021 system were averaged by calculating a consensus shape via the function *mShape* in geomorph v4.0.1 (Baken et al. 2021).

90-tip subclade-level tree details

We produced a subclade level tree of 90 tips composed of non-Passeriforme orders and Passeriforme families, only retaining the orders/families with 15 or more species (see Supplementary Tables S1-S2). Following Guillerme et al. (2023), we removed any orders represented by fewer than 15 tips in the MMB dataset, further decreasing the number of orders used in all analyses from 35 to 26. The excluded orders with associated species richnesses are: Leptosomiformes (1), Opisthocomiformes (1), Eurypygiformes (2), Cariamiformes (2), Mesitornithiformes (3), Phaethontiformes (3), Gaviiformes (5), Phoenicopteriformes (6), Coliiformes (6), Cathartiformes (7) (Table S3). Passeriformes, containing over half of all extant birds (Clements et al. 2021), were initially split into 141 family-level subclades, of which 77 were excluded (< 15 species per group; Table S4). Full lists of included and excluded non-passerine orders are available in Table S3 and for passerine families in Table S4. Summary statistics for each bird order are available in Table S1, containing information on order age, number of species in MMB, actual species richness, mean mass (kg) per order, and mean beak length (mm) per order. Ultimately, each of the main sampled clades was represented by more than 61% of recognized species (average 81.2%; Fig. 1b; Table. S3).

Per-subclade allometric and phylogenetic correction

We performed allometric and phylogenetic signal corrections for each of the 90 subclades. First, an ordinary least squares (OLS) was fit using landmarks as the response variable and (log) centroid size as predictor using the function *procD.lm* in geomorph v4.0.1 (using 10,000 permutations) (Table S5). Principal component analysis (PCA) was performed on these OLS residual landmarks, where all PCs were subjected to model fitting for best fitting Pagel's $\lambda$ using *fit_t_pl* in RPANDA v2.5 (Morlon et al. 2016). *fit_t_pl* is able to handle high-dimensional trait matrices via a special penalized likelihood approach using ridge regression and leave-one-out cross validation (PL-LOOCV) (Clavel et al. 2019). *fit_t_pl* was run per subclade using quadratic ridge penalty, null target matrix, and PL-LOOCV, extracting the phylogeny scaled to the estimated $\lambda$ parameter. Using best-fitting Pagel's $\lambda$ estimates (Table S6), we ran additional instances of *procD.lm* (10,000 permutations) using phylogenetic covariance matrices from the $\lambda$-

rescaled subclade phylogenies (essentially PGLS) (Table S7). We then compared the relative fit of OLS and PGLS models by calculating AIC via the function *model.comparison* in RRPP (Collyer and Adams 2018) (Table S8). Residual landmarks from the best fitting linear model (lowest AIC) were used in downstream analyses.

Gini coefficient demonstration figure

The following procedure describes analytical steps taken in the examples shown in Supplementary Fig. S1, but also describe the same steps using empirical data analyzed in the main text (AVONET discrete traits). This figure demonstrates the derivation of dietary ecological specialization via the Gini coefficient (GC), based on stochastic character mapping for a simulated generalist (**A**) and specialist (**B**) trait distribution. The following steps 1-4 apply to both panels A and B: **1.** A time-calibrated phylogeny along with a set of simulated discrete states per tip. For the purposes of this example, the phylogeny was simulated as a pure birth tree with 50 tips (via *pbtree* in phytools 2.5-2 (Revell 2024)) and the discrete states were randomly sampled with the Aquatic Predator niche arbitrarily chosen as the most common state, setting its sampling probability unevenly higher than the other eight states (using *sample* in base R). **2.** Via maximum likelihood model fitting, three modes of discrete character evolution were fitted: Equal Rates (ER), where transition rates between all states are equal (equivalent to the Mk model (Lewis 2001)); Symmetric Rates (SYM), where only forward and backward transition rates between each unique pair of traits are equal; All Rates Different (ARD) where forward and backwards rates are unequal for each unique pair of traits. After calculation of log-likelihood, model selection is performed by evaluating $AIC_c$ for each model. **3.** Based on the best fitting discrete evolutionary mode (i.e., lowest $AIC_c$ score), we performed stochastic character mapping (Bollback 2006) multiple times for the same tree. Inputs are the time-calibrated phylogeny, discrete traits, and best fitting evolutionary mode which are used in the *fastSimmap* function in the R package ratematrix 1.2.4 (Caetano and Harmon 2017). **4.** For each simulated character history, we calculate the total times spent in each discrete trait per branch of the phylogeny (i), then summed over each branch (ii). We then order the total times spent in each state in increasing order, calculate the cumulative share of each state (x) and the cumulative sum of each state, from which we calculate a cumulative proportion of each state (y). GC is typically calculated as the area between the curve after plotting the cumulative share of each state on the x-axis, and the cumulative proportion of each state on the y-axis, known as the Lorenz Curve (Lorenz 1905). A one-to-one line in the context of ecological states would represent the expectation from perfect ecological generalism. Here we calculate GC in an equivalent manner using the function *Gini* in the R package DescTools 0.99.6 (Signorell 2025).

Note on Morphospace Patterns of Recently Phylogenomically-Resolved Bird Clades

Several bird clades with depauperate morphospace representations (Fig. 1B) have recently gained phylogenetic resolution (Stiller et al. 2024). Principal findings include Mirandornithes (grebes, flamingos) having diverged from all other Neoavians about 67.4 Ma (pre-KPg boundary), and were recovered as sister to all other Neoavians. Assuming Mirandornithes have maintained aquatic habits since their origin, early members may have encountered abundant opportunities to expand within aquatic niches, as more derived aquatic Neoavian clades had not evolved yet. Although Elementaves (containing Aequornithes) were placed as sister to Telluraves (post-KPg divergence), the credible interval between the node uniting Elementaves and Telluraves and the crown age for Elementaves includes the KPg. Therefore, there may be a

connection in pre-KPg bird clades with aquatic habits and sparse morphospace representation. Perhaps these bird clades were more abundant throughout the Cenozoic but were outcompeted by later diverging aquatic bird clades (e.g. Charadriiformes, Gruiformes, Phaethontiformes), leading to morphospace culling by extinction and poor representation in the extant morphospace. A similar argument could be made for Paleognathae, which also have low morphospace density. As the earliest diverging modern bird "superorder", relatively higher abundance in the Cenozoic gave way to competition-induced extinction either with later diverging Neoavian clades, or more likely competition with other terrestrial competitors such as mammals, which had undergone an explosive adaptive radiation in the aftermath of the Kpg extinction (Halliday et al. 2016).
Low morphospace density and occupation of these clades may be driven by allometric effects to an extent, as these clades include relatively larger-bodied forms (e.g. ostriches, pelicans, flamingoes). Smaller-bodied birds have been shown to exhibit lower wing bone integration (Orkney et al. 2021), which may have facilitated not only wing shape diversification, but beak shape as well (assuming at least modest levels of integration with the beak). Paleognaths appear to be under significant allometric influence, albeit with moderate effect size ($p = 0.038$, $Z = 1.74$; Table S7), while Sphenisciformes (order of Aequornithes) are also under marginally insignificant allometric influence ($p = 0.076$, $Z = 1.46$; Table S7).

Discussion of Outlier Clades

Although not included in the PGLS analyses, the outlier subclades detected in our analyses represent potential cases of independent dynamics compared to the avian macroevolutionary norm. Between integration and ecological specialization, most of the outlier subclades are passerine families, with the exception of Musophagiformes (Fig. S8). Ptilorhynchidae, Paradisaeidae, Dicruridae, and Musophagiformes lie outside the top edges of the confidence ellipse. Ptilorhynchidae (Bowerbirds) and Paradisaeidae (Birds of Paradise) are most well-known for their elaborate plumage displays, proving to be model systems for sexual selection (Diamond 1986). In a review by Diamond (1986), it was noted that ptilorhynchids are fairly conserved in their bill morphology, while paradisaeids are quite variable. This may translate to lower bill disparity in ptilorhynchids and higher disparity in paradisaeids, which we can confirm by noting ptilorhynchids lie withing the CI ellipse, while paradisaeids lie outside the CI ellipse at least between disparity and ecological specialization (Fig. S10). The two passerine families are also rate-integration outliers, displaying especially high values of integration (Fig. S9). Although bill morphology may not be of greatest interest in Bowerbirds and Birds of Paradise compared to plumage and sexual selective traits, perhaps their elaborate displays are indirectly correlated with bill morphology, which may manifest in their overall levels of integration. Another outlier in terms of integration and dietary specialization is the passerine family includes Dicruridae (Drongos), whose members display elaborate plumage with forked tail feathers (Mayr and Vaurie 1948), further adding potential support to the speculation that high integration in bills occurs in birds with elaborate plumage structures.

Phylloscopidae are not only the passerine family with the highest level of integration and highest rate, but are also the largest outlier in these variables by an extreme extent (Fig. S9). This pattern appears to be robust to any possible effects of species richness and clade age. Assuming this is due to true biological signal, this family would represent an interesting case where the positive association between rates and integration as seen across most avian subclades is taken to an excessive extreme. Moreover, previous literature describing their morphology suggests limited morphological disparity (Tietze et al. 2015), which we corroborate with the plot between disparity and ecological specialization shows this clade displays a somewhat intermediate value of disparity. A significant disadvantage in attempting explanations for high rates, high integration, but low disparity in this particular family is the lack of literature describing their macroevolution or macroecology in the context of bill morphology.

Oriolidae (Old World Orioles), Mimidae (Mimid Thrushes), and Emberizidae (buntings) appear to be major outliers in terms of dietary specialization, occupying a small cluster in the far negative end of the Gini coefficients (Figs. S8-10), indicative of extreme dietary generalism. This pattern appears to be robust to corrections for clade age and species richness, although none of these families appear have the lowest species richness or significantly younger or older than other passerine families (Table S2). Literature on macroecological or macroevolutionary patterns in these families is extremely scant, severely limiting our interpretations regarding their potentially anomalously high levels of dietary generalism.

The orders Musophagiformes, Strisores, Ciconiiformes, Anseriformes, Psittaciformes, Accipitriformes, and the 'superorder' Paleognathae are outliers with respect to either clade age or species richness. Musophagiformes have limited discussion in the literature regarding these macroevolutionary variables, but their outlier status may be due to the presence of a casque along the beak in two species of *Musophaga* (Mayr 2018). Unusual structures may be biasing estimates, especially in an order with low total species richness (23 species, Table S2), but even lower sample in the MMB dataset (15, Table S2). Caprimulgiformes are an unusual order, represented by two major clusters in beak morphospace with Trochilidae dominating one end (long, slender, needle-like bills) and very broad-billed Bucerotiformes, Podargidae, Caprimulgidae, with a wide empty space lacking in intermediate forms between the two extremes (Fig. 2C). Such a wide gap is the likely source of high disparity and rates within the whole order, although elevated per-branch rate estimates have been documented in the broad-billed frogmouths, potoos, nightjars and along the basal branch leading to Trochilidae (hummingbirds) (Cooney et al. 2017).

Ciconiiformes (storks) display low rates, low disparity, and somewhat intermediate values of dietary specialization and integration (Fig. S8). Ciconiiformes are only marginally outside the confidence ellipses whenever they are retrieved as outliers, and also appear to lack extensive literature documenting their macroecology or macroevolution. Anseriformes (ducks, geese) display high rates, but somewhat intermediate values of integration and dietary specialization. Anseriformes are a mainly herbivorous radiation, with some lineages appearing to have evolved even greater specialization to herbivory compared to the anseriforme norm, at least five times (Olsen 2015). Broad links between bill morphology and herbivory, combined with multiple transitions to greater herbivore specialization, may have resulted in greater overall rates of bill morphology. Evidence of different selection regimes in terms of diet and foraging type in Anseriformes suggests links between diet and skull morphology exist (Chatterji et al. 2024). Accipitriformes (eagles, hawks) have previously been noted as unusual only in their relatively lower predicted mechanical advantage (MA) from regressions between beak shape and MA, but likely caused by use of talons for kill prey (Navalón et al. 2019). Psittaciformes in contrast display extreme MA, complemented by much slower gape velocity due to specializations on hardened food items (nuts, seeds) (Navalón et al. 2019). Diet may play a limited role in the cranial evolution of Psittaciformes, however, with the possibility that integration, phylogenetic signal, and allometry playing a much larger roles than diet (Bright et al. 2019). Perhaps the outlier status of Accipitriformes in these analyses results from their extreme lack of biomechanical efficiency, while in Psittaciformes their extreme enhancement of biomechanical efficiency drives their outlier status.

It is unfortunate that Paleognathae must be excluded from these analyses, as they represent the earliest diverging modern bird orders. Several factors may influence their outlier status. A combination of low species richness (59 sp.) and ancient divergence in the Late Cretaceous (68 Ma) result in unusually low rates (Table S3). Integration levels are among the lowest for all avian orders (Fig. 3; Table S3). Low integration may result from averaging of 5 paleognath orders into one superorder, with some orders within Paleognathae displaying variable levels of integration. Their outlier status may also result from lifestyle and large mass, being primarily flightless ground-dwelling birds and displaying large mass disparity represented by the largest modern birds (Struthioniformes: ostriches, emus, rheas) and the smallest terrestrial birds (kiwis).

**Fig. S1**

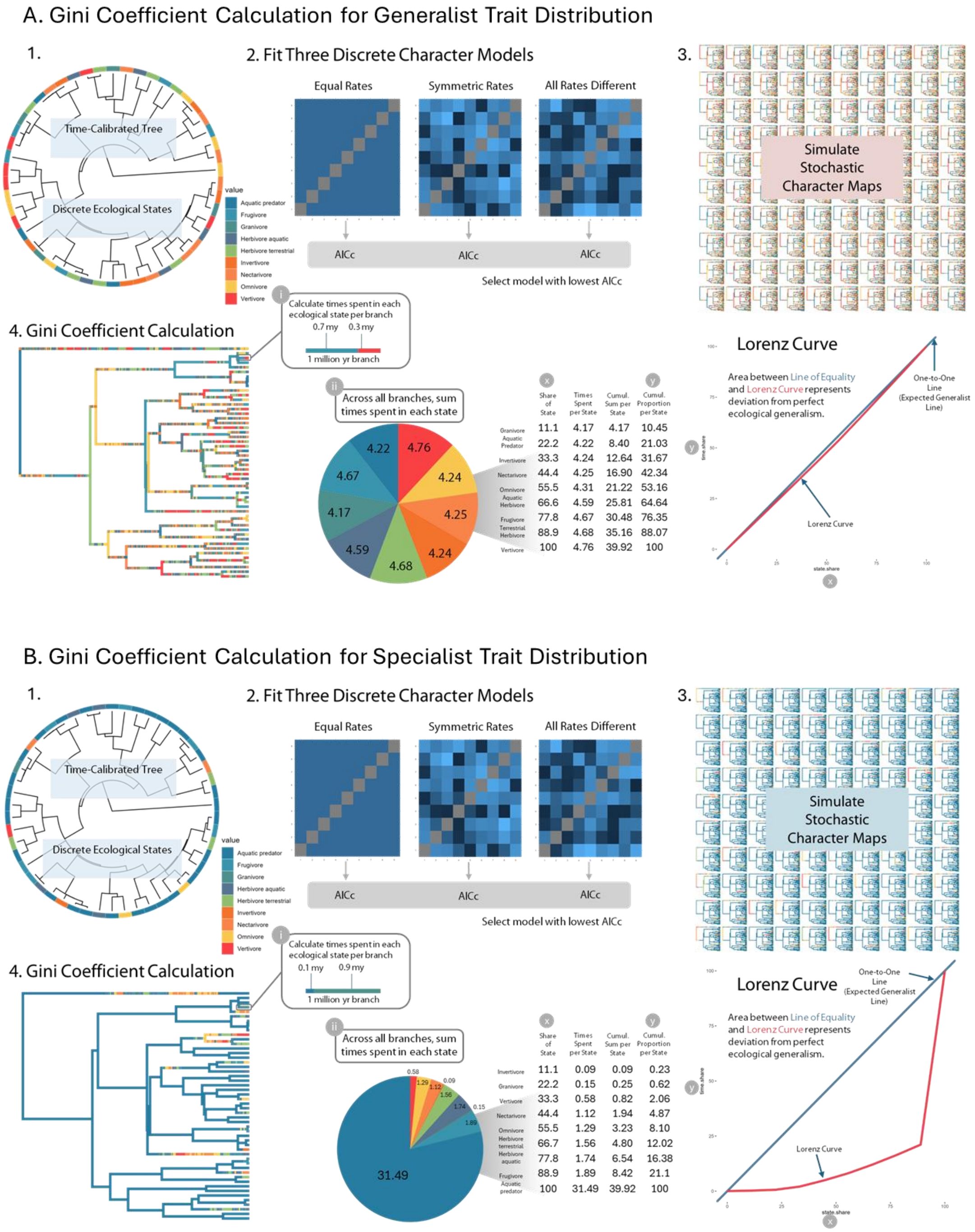

Dietary ecological specialization calculated by the Gini coefficient (GC), based on stochastic character mapping for simulated **A**) generalist and (**B**) specialist trait distributions. The following steps 1-4 apply to both panels A and B: **1.** Generate time-calibrated phylogeny with a set of discrete states per tip. **2.** Fit three modes of discrete character evolution: Equal Rates, Symmetric Rates, All Rates Different. **3.** Perform stochastic character mapping (Bollback 2006) from the best fitting model according to lowest AICc (Akaike Information Criterion corrected) value (Step 2) multiple times (1000 iterations). **4.** For each stochastic character map, calculate total times spent in each discrete trait per branch of the phylogeny (i), then sum over each branch (ii). Order total times spent in each state in increasing order, calculate the cumulative share of each state (x) and the cumulative sum of each state, from which to calculate a cumulative proportion of each state (y). GC is the area between the curve after plotting the cumulative share of each state on the x-axis, and the cumulative proportion of each state on the y-axis, known as the Lorenz Curve. See Supplementary Section Id for full methods description.

**Fig. S2.**

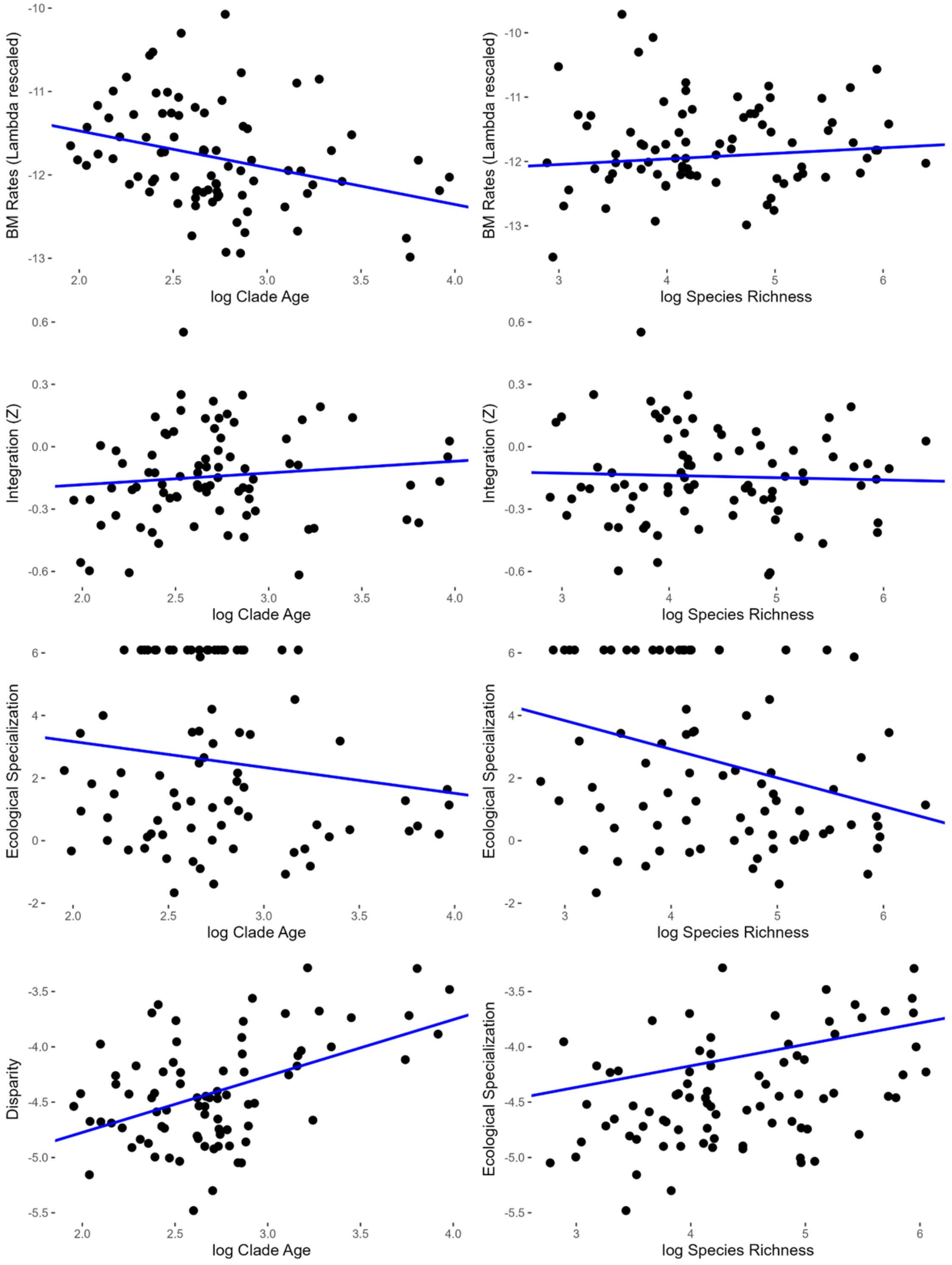

Species Richness and Clade Age PGLS regressions (with outliers excluded).

**Fig. S3.**

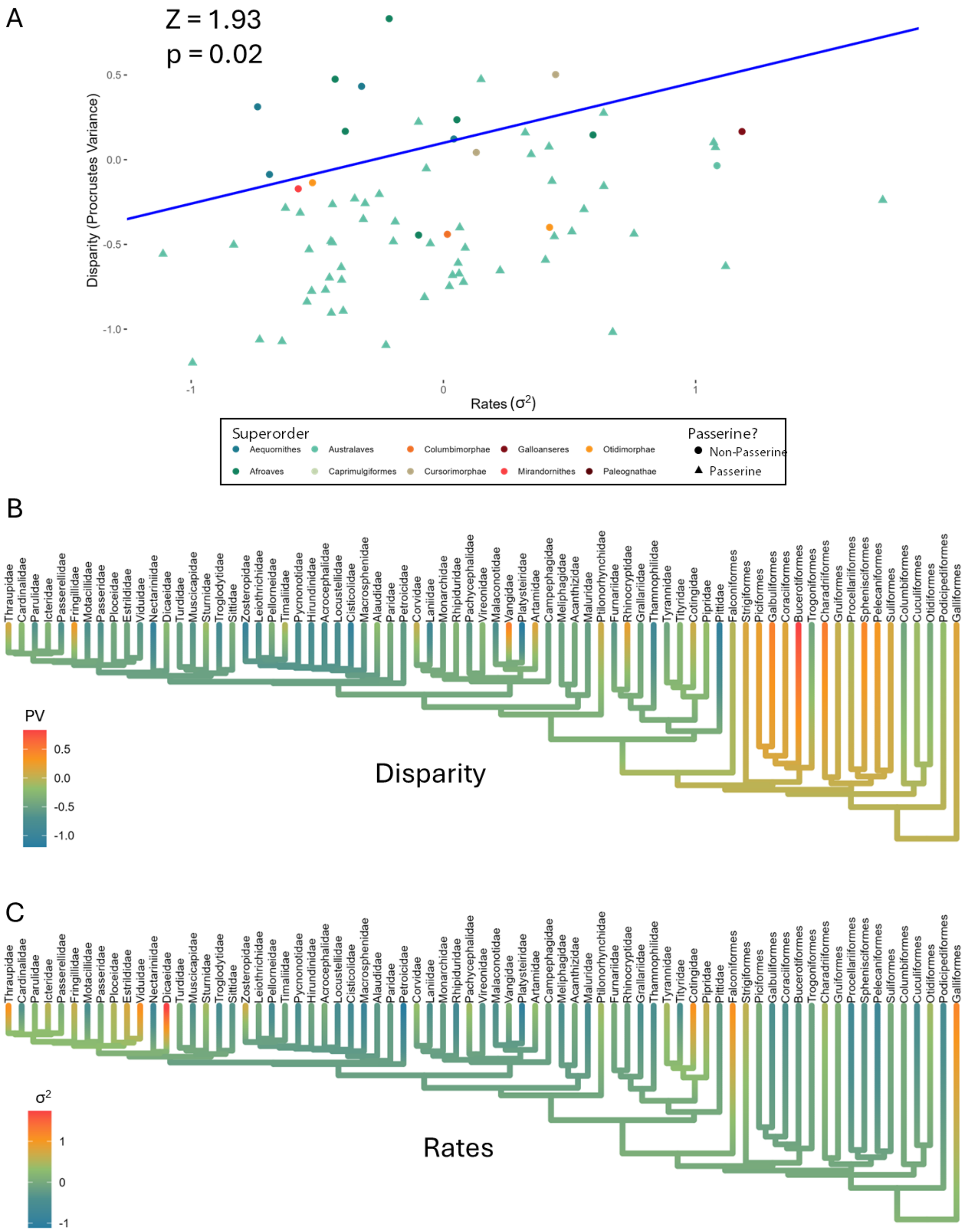

Relationships among morphological disparity as Procrustes variance (PV) and evolutionary rates ($\sigma^2$). Only non-outlier subclades are shown, and all values pertain to clade age and/or species richness PGLS residuals. A. PGLS of disparity regressed onto rates. B. The evolution of disparity mapped on the subclade-level phylogeny (according to a standard Brownian motion character). C. The evolution of rates mapped on the subclade-level phylogeny (according to a standard Brownian motion character). Made with coding solution by Claude Opus 4.6.

**Fig. S4**

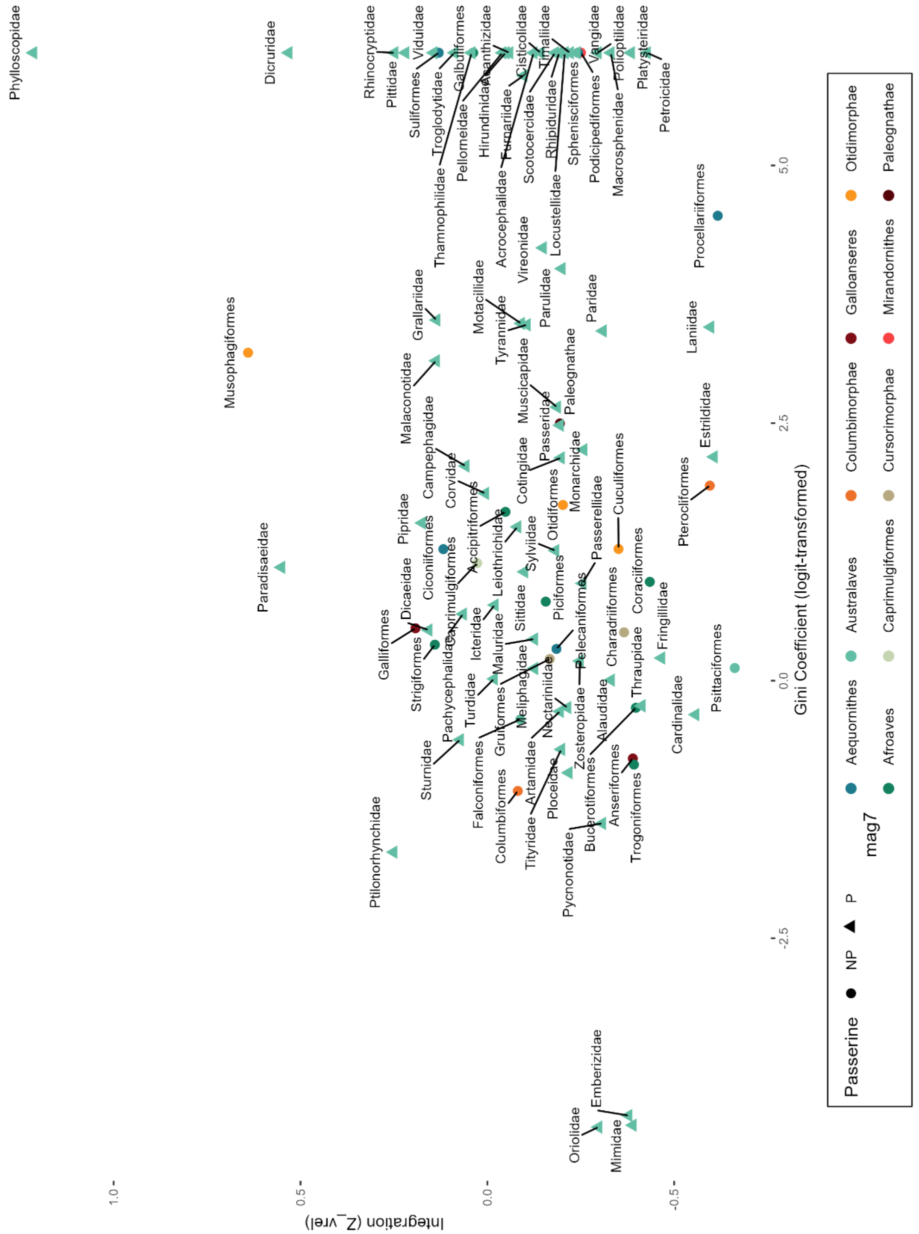

Integration ($Z_{Vrel}$) on ecological specialization (logit Gini), non-residual values.

**Fig. S5**

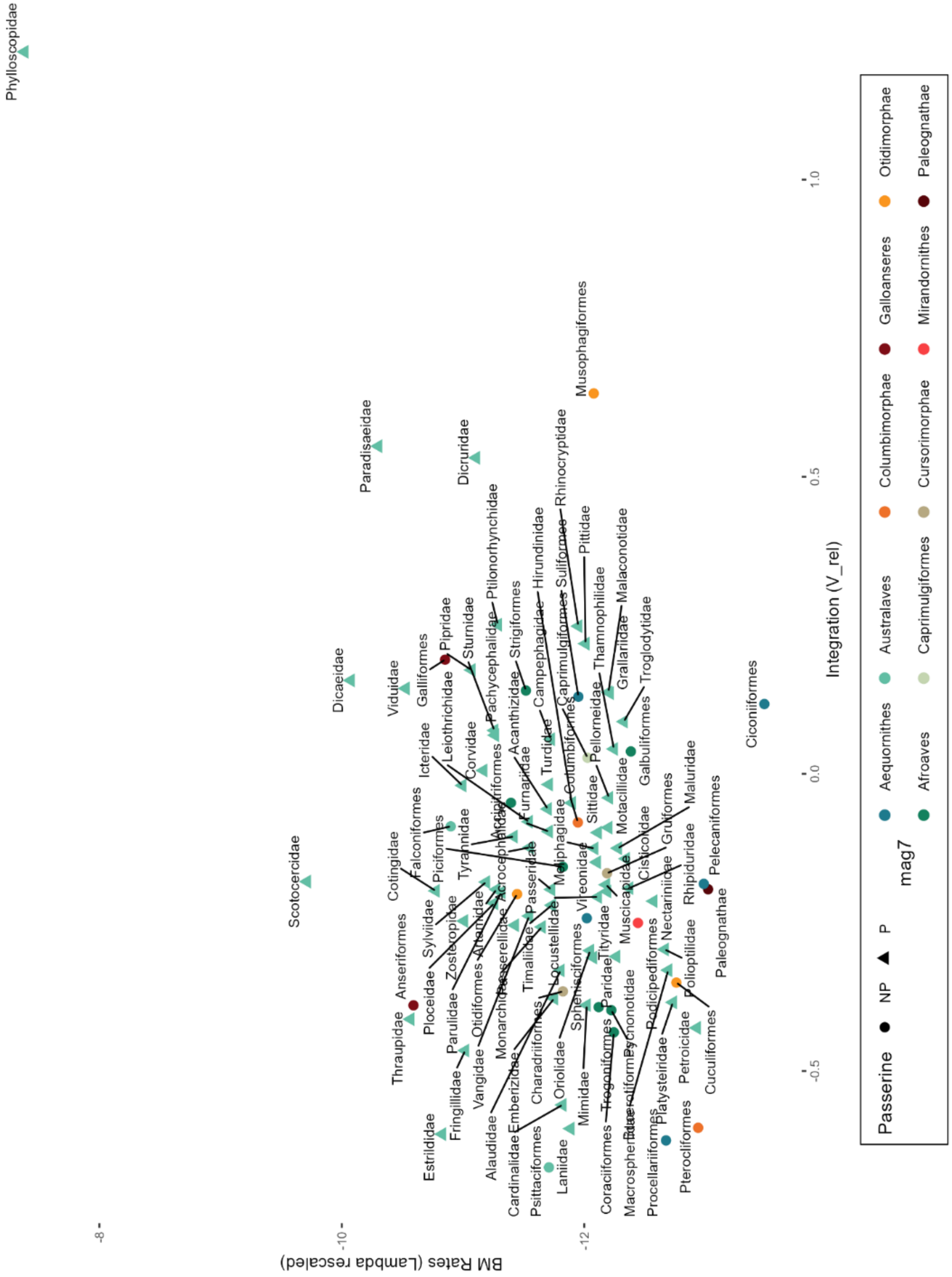

Rates on Integration, non-residual values.

**Fig. S6**

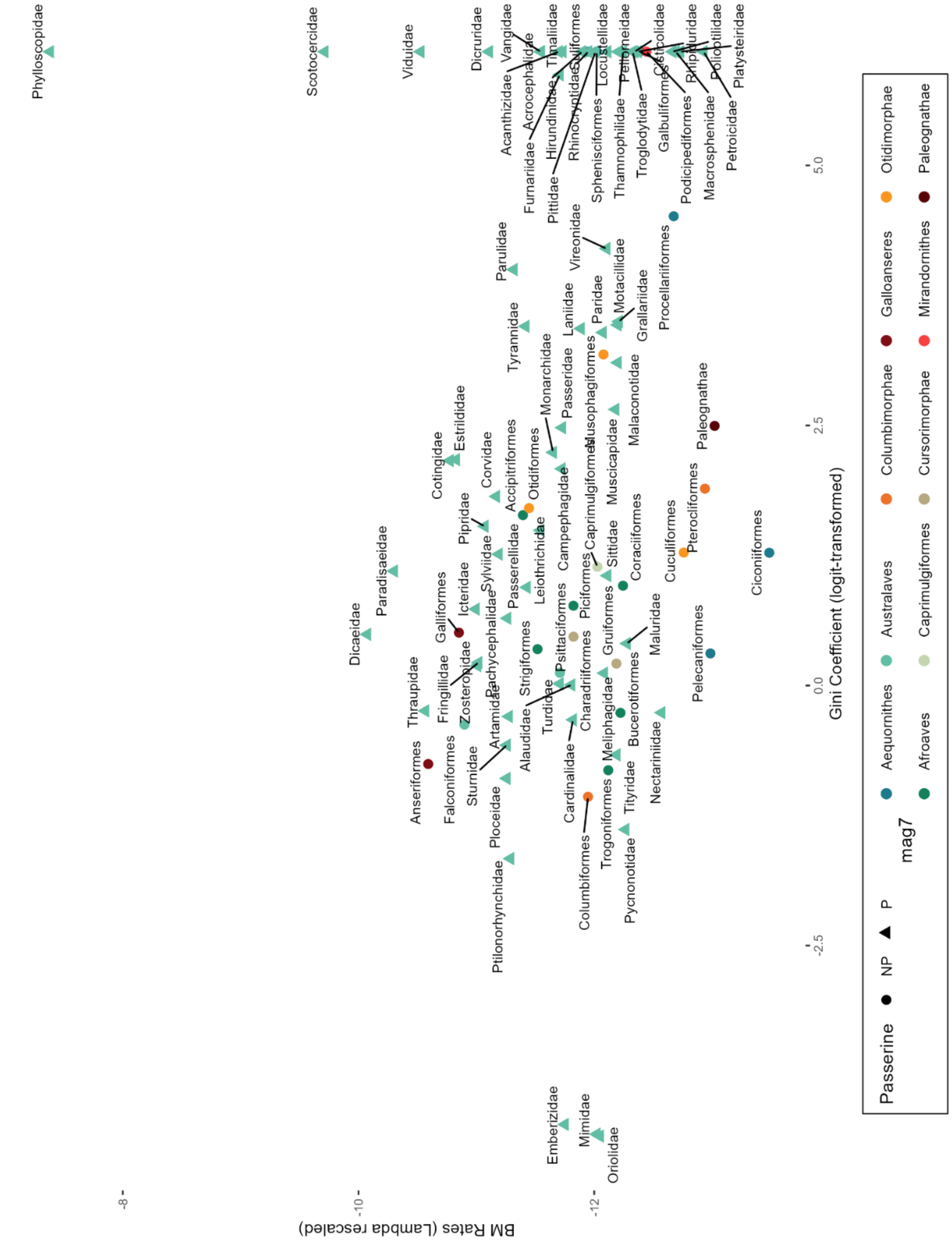

Rates on ecological specialization, non-residual values.

**Fig. S7**

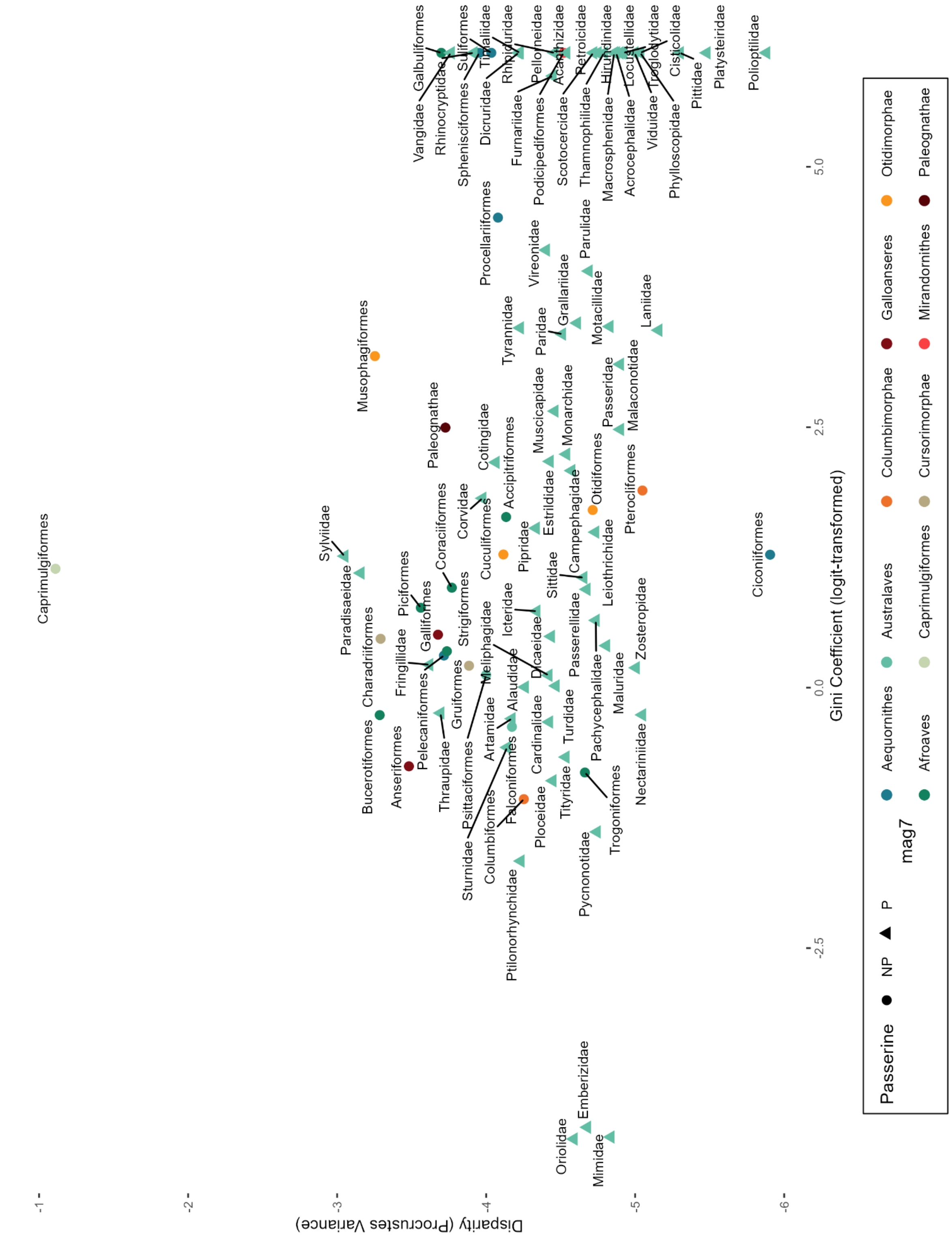

Disparity on ecological specialization, non-residual values.

**Fig. S8**

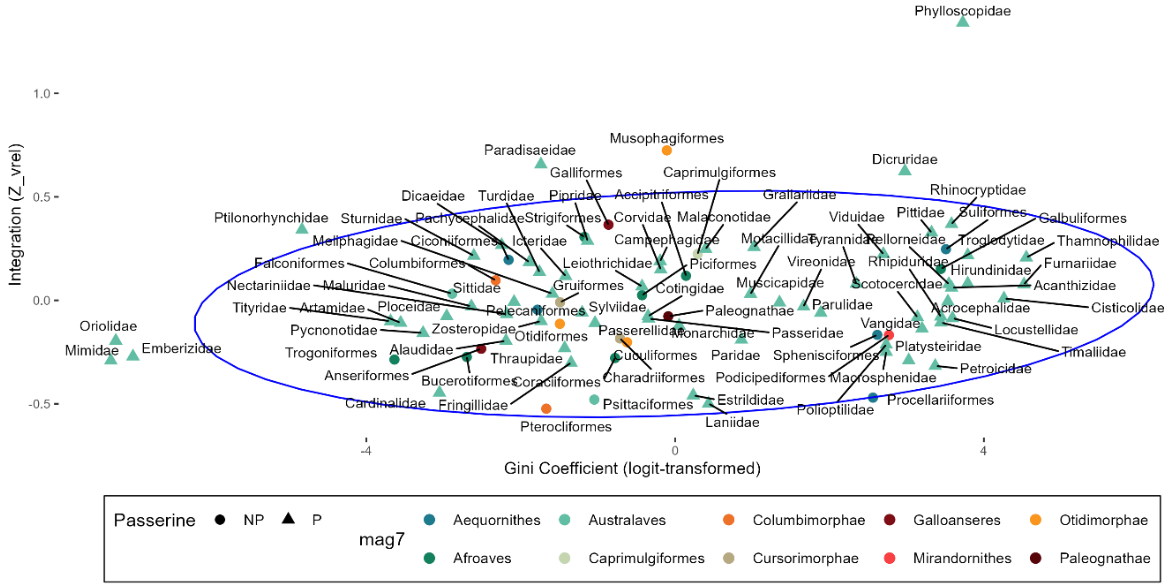

Integration vs Ecological Specialization (residuals via clade age and/or species richness). 95% Confidence Interval ellipse displayed.

**Fig. S9**

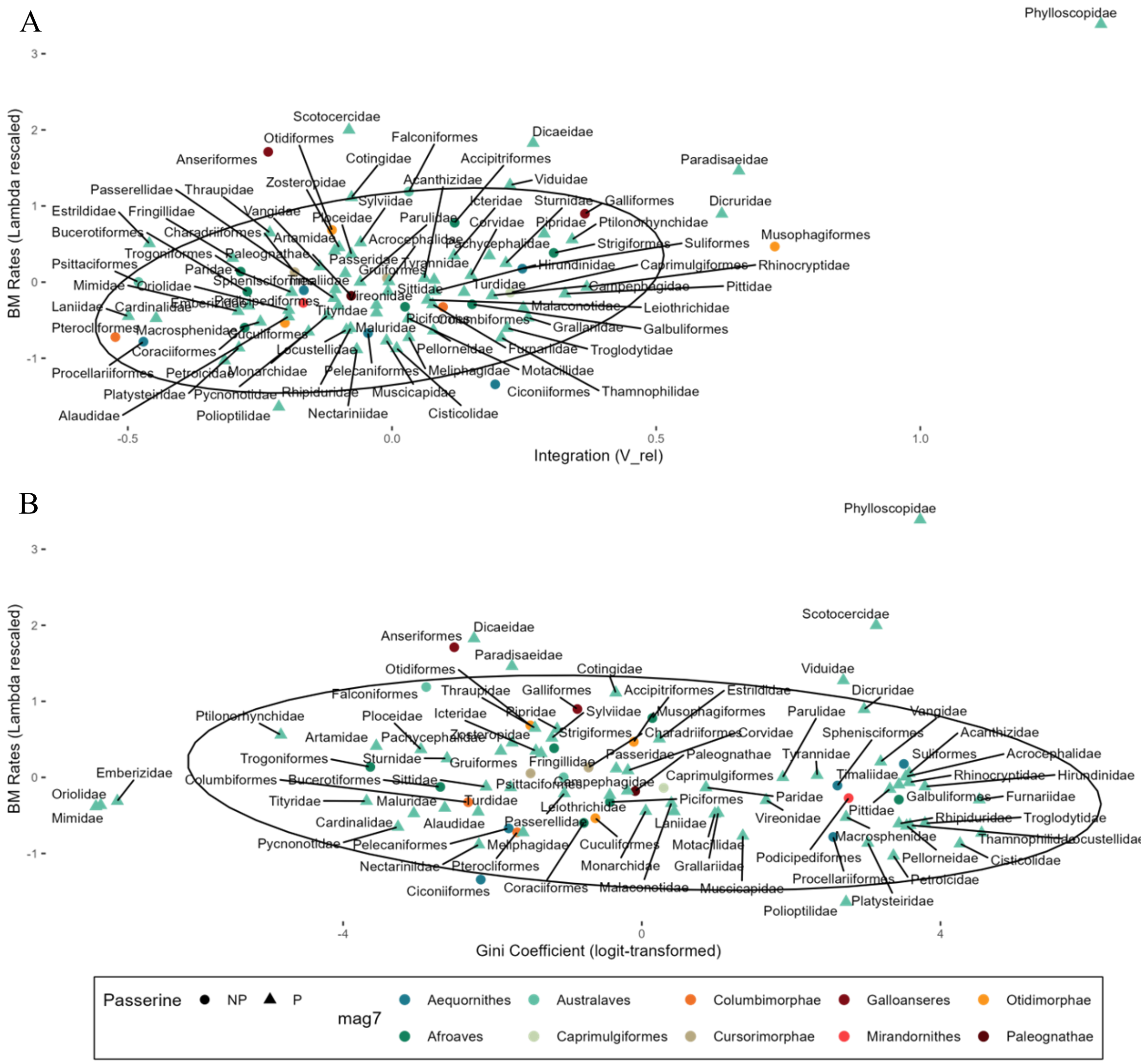

A. Rates on Integration. B. Rates on GC (residuals via clade age and/or species richness). 95% Confidence Interval ellipses displayed.

**Fig. S10.**

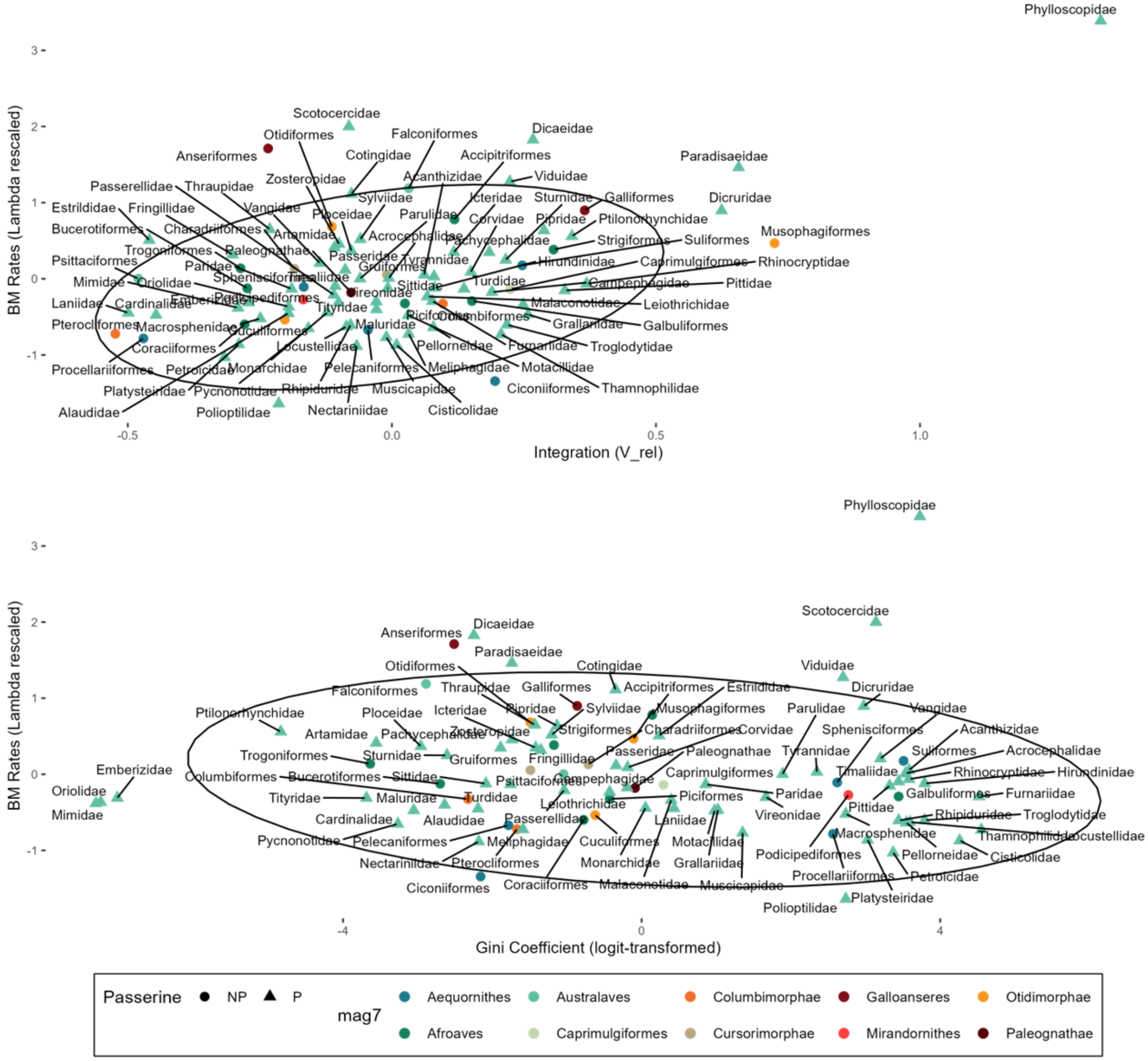

A. Rates on Integration. B. Rates on GC (residuals via clade age and/or species richness). 95% Confidence Interval ellipses displayed.

**Fig. S11.**

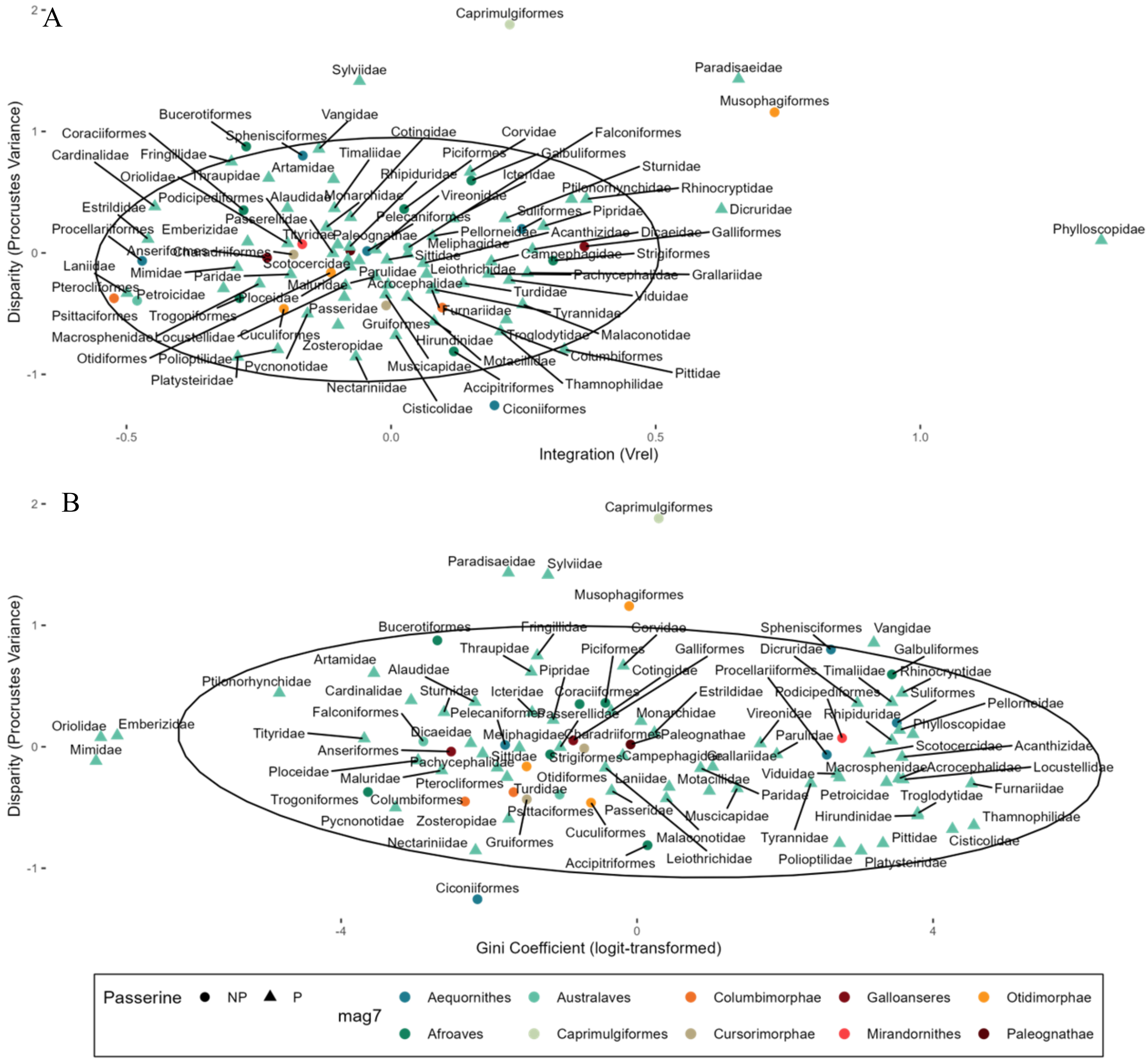

Disparity on Integration and GC (residuals via clade age and/or species richness). 95% Confidence Interval ellipses displayed.

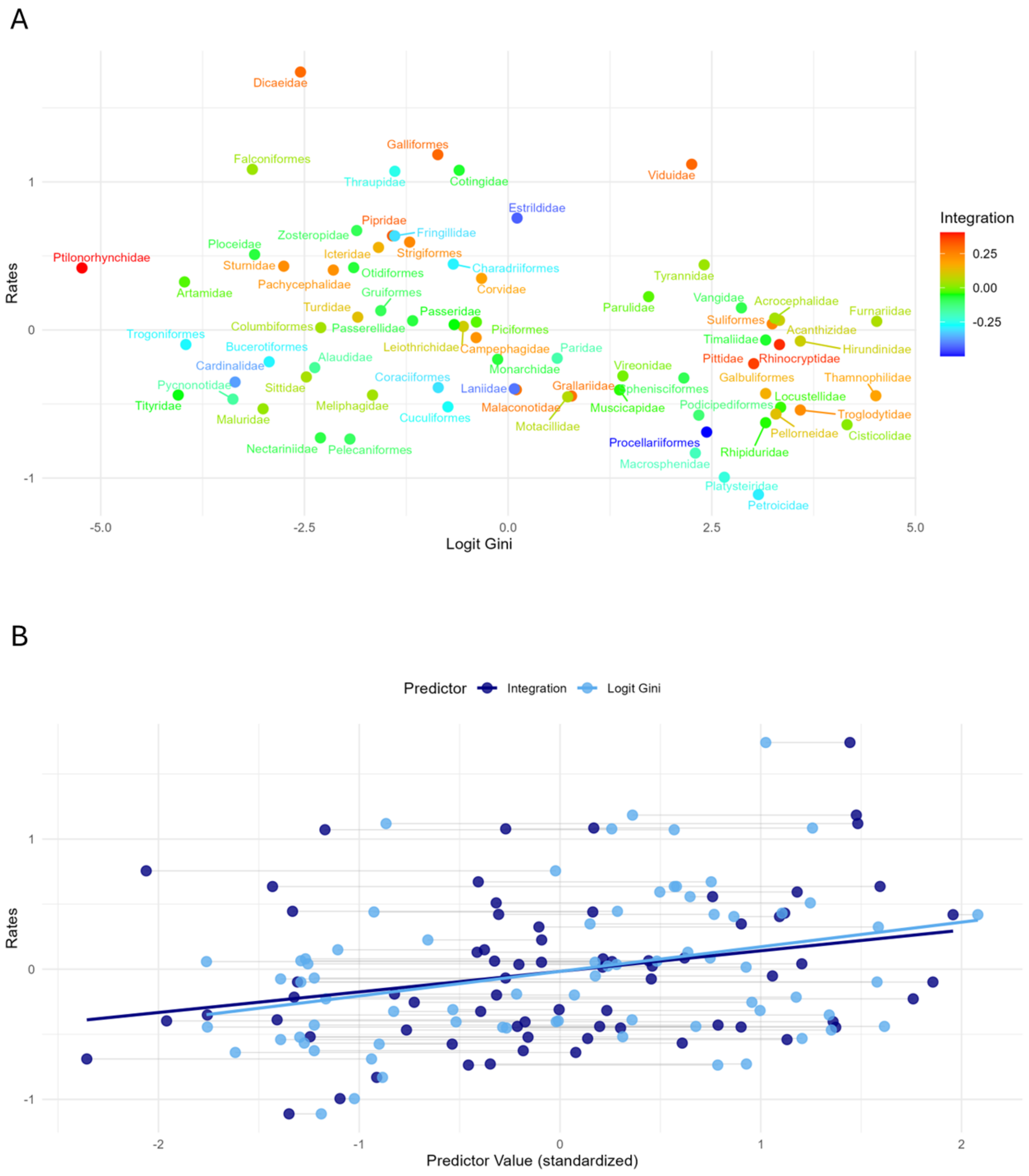

**Fig. S12**

Rates on (logit) Gini Coefficient, and their interaction with integration. Only residual values from clade age and/or species richness PGLS models were used, with outlier subclades excluded (based on CA and/or SR PGLS). A. Rates versus (logit) GC, with points colored according to magnitude of integration. B. Rates versus standardized (Z-transformation) of both integration and ecological specialization, superimposed. Importantly, logit Gini values were “inverted” (sign changed) such that greater values now reflect greater generalism, more diverse trophic histories, or greater number of discrete state changes. Horizontal lines connect integration and logit-GC scores of the same subclades.

**Table S1.**
List of included and excluded bird orders, based on whether each order contains greater than 15 taxa.

| Order | Species Number | Excluded? |
|---|---|---|
| Paleognathae | 59 | No |
| Anseriformes | 178 | No |
| Galliformes | 298 | No |
| Phoenicopteriformes | 6 | Yes |
| Podicipediformes | 22 | No |
| Columbiformes | 348 | No |
| Mesitornithiformes | 3 | Yes |
| Pterocliformes | 16 | No |
| Otidiformes | 26 | No |
| Musophagiformes | 23 | No |
| Cuculiformes | 147 | No |
| Caprimulgiformes | 599 | No |
| Gruiformes | 192 | No |
| Charadriiformes | 383 | No |
| Eurypygiformes | 2 | Yes |
| Phaethontiformes | 3 | Yes |
| Gaviiformes | 5 | Yes |
| Sphenisciformes | 18 | No |
| Procellariiformes | 138 | No |
| Ciconiiformes | 19 | No |
| Suliformes | 59 | No |
| Pelecaniformes | 114 | No |
| Cathartiformes | 7 | Yes |
| Accipitriformes | 252 | No |
| Strigiformes | 243 | No |
| Coliiformes | 6 | Yes |
| Leptosomiformes | 1 | Yes |
| Trogoniformes | 43 | No |
| Bucerotiformes | 72 | No |
| Coraciiformes | 183 | No |
| Galbuliformes | 54 | No |
| Piciformes | 377 | No |
| Cariamiformes | 2 | Yes |
| Falconiformes | 65 | No |
| Psittaciformes | 390 | No |
| Passeriformes | 6469 | No |

**Table S2.**
List of included and excluded Passeriforme families, based on whether the family contains greater than 15 taxa.

| Family | Species Number | Excluded? |
|---|---|---|
| Acanthisittidae | 4 | Yes |
| Calyptomenidae | 6 | Yes |
| Eurylaimidae | 9 | Yes |
| Sapayoidae | 1 | Yes |
| Philepittidae | 4 | Yes |
| Pittidae | 46 | No |
| Thamnophilidae | 237 | No |
| Melanopareiidae | 4 | Yes |
| Conopophagidae | 11 | Yes |
| Grallariidae | 68 | No |
| Rhinocryptidae | 65 | No |
| Formicariidae | 12 | Yes |
| Furnariidae | 306 | No |
| Pipridae | 53 | No |
| Cotingidae | 65 | No |
| Tityridae | 33 | No |
| Oxyruncidae | 7 | Yes |
| Tyrannidae | 425 | No |
| Menuridae | 2 | Yes |
| Atrichornithidae | 2 | Yes |
| Ptilonorhynchidae | 27 | No |
| Climacteridae | 7 | Yes |
| Maluridae | 32 | No |
| Meliphagidae | 190 | No |
| Dasyornithidae | 3 | Yes |
| Pardalotidae | 4 | Yes |
| Acanthizidae | 65 | No |
| Pomatostomidae | 5 | Yes |
| Orthonychidae | 3 | Yes |
| Cinclosomatidae | 12 | Yes |
| Campephagidae | 89 | No |
| Mohouidae | 3 | Yes |
| Neosittidae | 3 | Yes |
| Psophodidae | 5 | Yes |
| Eulacestomatidae | 1 | Yes |
| Oreoicidae | 3 | Yes |
| Falcunculidae | 1 | Yes |
| Paramythiidae | 2 | Yes |
| Vireonidae | 63 | No |
| Pachycephalidae | 63 | No |

| | | |
|---|---|---|
| Oriolidae | 38 | No |
| Machaerirhynchidae | 2 | Yes |
| Artamidae | 24 | No |
| Rhagologidae | 1 | Yes |
| Platysteiridae | 31 | No |
| Vangidae | 39 | No |
| Pityriasidae | 1 | Yes |
| Aegithinidae | 4 | Yes |
| Malaconotidae | 50 | No |
| Rhipiduridae | 54 | No |
| Dicruridae | 29 | No |
| Paradisaeidae | 42 | No |
| Ifritidae | 1 | Yes |
| Monarchidae | 100 | No |
| Corcoracidae | 2 | Yes |
| Melampittidae | 2 | Yes |
| Laniidae | 35 | No |
| Corvidae | 128 | No |
| Cnemophilidae | 3 | Yes |
| Melanocharitidae | 11 | Yes |
| Callaeidae | 5 | Yes |
| Notiomystidae | 1 | Yes |
| Petroicidae | 49 | No |
| Picathartidae | 2 | Yes |
| Chaetopidae | 2 | Yes |
| Eupetidae | 1 | Yes |
| Hyliotidae | 4 | Yes |
| Stenostiridae | 9 | Yes |
| Paridae | 63 | No |
| Remizidae | 11 | Yes |
| Alaudidae | 99 | No |
| Panuridae | 1 | Yes |
| Nicatoridae | 3 | Yes |
| Macrosphenidae | 18 | No |
| Acrocephalidae | 62 | No |
| Hyliidae | 2 | Yes |
| Cisticolidae | 161 | No |
| Locustellidae | 66 | No |
| Donacobiidae | 1 | Yes |
| Bernieridae | 11 | Yes |
| Pnoepygidae | 5 | Yes |
| Hirundinidae | 86 | No |
| Pycnonotidae | 151 | No |
| Phylloscopidae | 79 | No |
| Scotocercidae | 36 | No |

| | | |
|---|---|---|
| Aegithalidae | 11 | Yes |
| Sylviidae | 69 | No |
| Zosteropidae | 142 | No |
| Timaliidae | 54 | No |
| Pellorneidae | 62 | No |
| Leiothrichidae | 143 | No |
| Regulidae | 6 | Yes |
| Tichodromidae | 1 | Yes |
| Sittidae | 28 | No |
| Certhiidae | 11 | Yes |
| Polioptilidae | 21 | No |
| Troglodytidae | 86 | No |
| Elachuridae | 1 | Yes |
| Cinclidae | 5 | Yes |
| Buphagidae | 2 | Yes |
| Sturnidae | 123 | No |
| Mimidae | 34 | No |
| Turdidae | 175 | No |
| Muscicapidae | 326 | No |
| Bombycillidae | 3 | Yes |
| Mohoidae | 5 | Yes |
| Ptiliogonatidae | 4 | Yes |
| Dulidae | 1 | Yes |
| Hylocitreidae | 1 | Yes |
| Hypocoliidae | 1 | Yes |
| Promeropidae | 2 | Yes |
| Modulatricidae | 3 | Yes |
| Dicaeidae | 48 | No |
| Nectariniidae | 143 | No |
| Irenidae | 2 | Yes |
| Chloropseidae | 11 | Yes |
| Peucedramidae | 1 | Yes |
| Urocynchramidae | 1 | Yes |
| Ploceidae | 118 | No |
| Estrildidae | 140 | No |
| Viduidae | 20 | No |
| Prunellidae | 13 | Yes |
| Passeridae | 43 | No |
| Motacillidae | 67 | No |
| Fringillidae | 229 | No |
| Calcariidae | 6 | Yes |
| Rhodinocichlidae | 1 | Yes |
| Emberizidae | 44 | No |
| Passerellidae | 132 | No |
| Calyptophilidae | 2 | Yes |

| | | |
|---|---|---|
| Phaenicophilidae | 4 | Yes |
| Nesospingidae | 1 | Yes |
| Spindalidae | 4 | Yes |
| Zeledoniidae | 1 | Yes |
| Teretistridae | 2 | Yes |
| Icteriidae | 1 | Yes |
| Icteridae | 105 | No |
| Parulidae | 111 | No |
| Mitrospingidae | 4 | Yes |
| Cardinalidae | 49 | No |
| Thraupidae | 381 | No |

**Table S3.**
Summary statistics of sampled bird orders. including clade age (crown), species richness (SR) in both the MMB dataset and actual SR, percent coverage, or percent of MMB species out of actual SR, mean mass (kg), standard deviation (SD) of mass (kg), mean beak length (mm), SD of beak length.

| | **Clade Age** | **SR (MMB)** | **SR (Actual)** | **Percent Coverag e** | **Mass (mean)** | **Mass (SD)** | **Beak Length (mean)** | **Beak Length (SD)** |
|---|---|---|---|---|---|---|---|---|
| Accipitriformes | 52.53 | 227 | 252 | 90.08 | 1.27 | 1.66 | 39.27 | 1.27 |
| Anseriformes | 53.53 | 154 | 178 | 86.52 | 1.49 | 1.62 | 48.97 | 1.49 |
| Bucerotiformes | 24.9 | 65 | 72 | 90.28 | 0.82 | 0.9 | 117.86 | 0.82 |
| Caprimulgiformes | 53.02 | 367 | 599 | 61.27 | 0.03 | 0.05 | 19.61 | 0.03 |
| Charadriiformes | 44.95 | 354 | 383 | 92.43 | 0.27 | 0.3 | 44.15 | 0.27 |
| Ciconiiformes | 16.69 | 19 | 19 | 100 | 3.67 | 2.06 | 226.86 | 3.67 |
| Columbiformes | 22.49 | 265 | 348 | 76.15 | 0.26 | 0.26 | 22.38 | 0.26 |
| Coraciiformes | 17.6 | 149 | 183 | 81.42 | 0.07 | 0.06 | 42.86 | 0.07 |
| Cuculiformes | 42.18 | 129 | 147 | 87.76 | 0.15 | 0.14 | 32.12 | 0.15 |
| Falconiformes | 23.53 | 61 | 65 | 93.85 | 0.37 | 0.31 | 24.68 | 0.37 |
| Galbuliformes | 22.09 | 43 | 54 | 79.63 | 0.04 | 0.02 | 36.36 | 0.04 |
| Galliformes | 26.51 | 263 | 298 | 88.26 | 0.9 | 0.88 | 29.06 | 0.9 |
| Gruiformes | 50.32 | 136 | 192 | 70.83 | 0.78 | 1.56 | 44.3 | 0.78 |
| Musophagiformes | 29.95 | 22 | 23 | 95.65 | 0.31 | 0.16 | 27.33 | 0.31 |
| Otidiformes | 18.09 | 25 | 26 | 96.15 | 2.57 | 2.35 | 53.24 | 2.57 |
| Paleognathae | 68.43 | 51 | 59 | 86.44 | 7.84 | 22.21 | 46.37 | 7.84 |
| Passeriformes | 26.05 | 5155 | 6469 | 79.69 | 0.04 | 0.06 | 18.61 | 0.04 |
| Pelecaniformes | 43.02 | 102 | 114 | 89.47 | 1.32 | 1.66 | 129.33 | 1.32 |
| Piciformes | 18.49 | 340 | 377 | 90.19 | 0.09 | 0.1 | 32.34 | 0.09 |
| Podicipediformes | 18.09 | 18 | 22 | 81.82 | 0.55 | 0.41 | 38.2 | 0.55 |
| Procellariiformes | 23.63 | 109 | 138 | 78.99 | 0.76 | 1.35 | 42.26 | 0.76 |
| Psittaciformes | 28.29 | 340 | 390 | 87.18 | 0.19 | 0.23 | 26.18 | 0.19 |
| Pterocliformes | 17.44 | 16 | 16 | 100 | 0.24 | 0.07 | 18.19 | 0.24 |
| Sphenisciformes | 12.29 | 17 | 18 | 94.44 | 5.91 | 7.27 | 64.12 | 5.91 |
| Strigiformes | 31.51 | 71 | 243 | 29.22 | 0.35 | 0.49 | 27.12 | 0.35 |
| Suliformes | 24.05 | 52 | 59 | 88.14 | 1.7 | 0.62 | 73.73 | 1.7 |
| Trogoniformes | 25.62 | 40 | 43 | 93.02 | 0.09 | 0.04 | 21.11 | 0.09 |

**Table S4.**
Summary statistics per Passeriforme family, including clade age (crown), species richness (SR) in both the MMB dataset and actual SR, percent coverage, or percent of MMB species out of actual SR, mean mass (kg), standard deviation (SD) of mass(kg), mean beak length, SD of beak length.

| Clade | Clade Age | SR (MMB) | SR (Actual) | Percent Coverage | Mass (mean) | Mass (SD) | Beak Length (Mean) | Beak Length (SD) |
|---|---|---|---|---|---|---|---|---|
| Acanthizidae | 14.32 | 54 | 65 | 83.08 | 0.011 | 0.005 | 12.719 | 2.25 |
| Acrocephalidae | 10.55 | 50 | 62 | 80.65 | 0.018 | 0.009 | 19.029 | 5.359 |
| Alaudidae | 8.84 | 86 | 99 | 86.87 | 0.03 | 0.012 | 15.86 | 3.865 |
| Artamidae | 9.87 | 22 | 24 | 91.67 | 0.112 | 0.113 | 36.033 | 16.064 |
| Campephagidae | 11.64 | 73 | 89 | 82.02 | 0.055 | 0.035 | 22.957 | 6.498 |
| Cardinalidae | 7.32 | 45 | 49 | 91.84 | 0.032 | 0.015 | 17.261 | 3.644 |
| Cisticolidae | 12.49 | 138 | 161 | 85.71 | 0.011 | 0.004 | 13.571 | 2.132 |
| Corvidae | 8.15 | 110 | 128 | 85.94 | 0.249 | 0.196 | 42.971 | 12.842 |
| Cotingidae | 17.51 | 57 | 65 | 87.69 | 0.109 | 0.089 | 23.108 | 8.157 |
| Dicaeidae | 16.08 | 42 | 48 | 87.5 | 0.008 | 0.001 | 11.002 | 1.078 |
| Dicruridae | 15.8 | 17 | 29 | 58.62 | 0.055 | 0.023 | 28.652 | 5.807 |
| Emberizidae | 8.17 | 41 | 44 | 93.18 | 0.02 | 0.006 | 13.034 | 1.624 |
| Estrildidae | 9.5 | 130 | 140 | 92.86 | 0.012 | 0.004 | 11.466 | 1.758 |
| Fringillidae | 11.13 | 187 | 229 | 81.66 | 0.022 | 0.013 | 13.879 | 4.175 |
| Furnariidae | 14.38 | 239 | 306 | 78.1 | 0.031 | 0.021 | 21.922 | 10.338 |
| Grallariidae | 14.3 | 28 | 68 | 41.18 | 0.06 | 0.045 | 23.271 | 4.916 |
| Hirundinidae | 16.34 | 76 | 86 | 88.37 | 0.019 | 0.01 | 11.01 | 2.267 |
| Icteridae | 8.85 | 94 | 105 | 89.52 | 0.078 | 0.073 | 28.889 | 11.324 |
| Laniidae | 7.68 | 31 | 35 | 88.57 | 0.047 | 0.021 | 21.014 | 2.708 |
| Leiothrichidae | 9.16 | 131 | 143 | 91.61 | 0.062 | 0.028 | 22.499 | 4.842 |
| Locustellidae | 9.65 | 46 | 66 | 69.7 | 0.023 | 0.016 | 16.489 | 4.083 |
| Macrosphenidae | 17.85 | 17 | 18 | 94.44 | 0.014 | 0.008 | 15.478 | 3.335 |
| Malaconotidae | 15.39 | 44 | 50 | 88 | 0.046 | 0.017 | 24.502 | 4.09 |
| Maluridae | 13.72 | 21 | 32 | 65.63 | 0.014 | 0.007 | 14.103 | 2.425 |
| Meliphagidae | 10.91 | 155 | 190 | 81.58 | 0.036 | 0.036 | 24.042 | 8.371 |
| Mimidae | 10.1 | 34 | 34 | 100 | 0.062 | 0.013 | 28.441 | 6.173 |
| Monarchidae | 7.06 | 26 | 100 | 26 | 0.02 | 0.011 | 18.401 | 3.224 |
| Motacillidae | 13.8 | 59 | 67 | 88.06 | 0.025 | 0.009 | 16.331 | 2.105 |
| Muscicapidae | 14.68 | 281 | 326 | 86.2 | 0.024 | 0.017 | 16.482 | 3.86 |
| Nectariniidae | 17.11 | 123 | 143 | 86.01 | 0.011 | 0.006 | 22.897 | 7.835 |
| Oriolidae | 11.03 | 32 | 38 | 84.21 | 0.077 | 0.023 | 29.528 | 4.526 |
| Pachycephalidae | 11.51 | 36 | 63 | 57.14 | 0.033 | 0.014 | 19.325 | 3.645 |
| Paradisaeidae | 12.71 | 39 | 42 | 92.86 | 0.166 | 0.069 | 41.44 | 15.184 |
| Paridae | 18.7 | 54 | 63 | 85.71 | 0.015 | 0.006 | 11.417 | 2.285 |
| Parulidae | 8.65 | 89 | 111 | 80.18 | 0.011 | 0.003 | 13.469 | 1.653 |

| | | | | | | | |
|---|---|---|---|---|---|---|---|
| Passerellidae | 7.7 | 109 | 132 | 82.58 | 0.028 | 0.01 | 15.356 | 2.202 |
| Passeridae | 14.3 | 40 | 43 | 93.02 | 0.025 | 0.007 | 13.57 | 1.797 |
| Pellorneidae | 10.74 | 54 | 62 | 87.1 | 0.026 | 0.011 | 18.6 | 5.394 |
| Petroicidae | 16.14 | 35 | 49 | 71.43 | 0.022 | 0.011 | 15.476 | 3.168 |
| Phylloscopidae | 3.43 | 64 | 79 | 81.01 | 0.008 | 0.002 | 11.784 | 1.635 |
| Pipridae | 12.54 | 48 | 53 | 90.57 | 0.014 | 0.005 | 11.96 | 1.632 |
| Pittidae | 14.92 | 32 | 46 | 69.57 | 0.084 | 0.031 | 25.641 | 3.94 |
| Platysteiridae | 13.46 | 24 | 31 | 77.42 | 0.012 | 0.004 | 15.248 | 1.57 |
| Ploceidae | 14.39 | 110 | 118 | 93.22 | 0.028 | 0.012 | 17.903 | 2.829 |
| Polioptilidae | 4.06 | 10 | 21 | 47.62 | 0.007 | 0.002 | 13.71 | 4.391 |
| Ptilonorhynchidae | 12.55 | 17 | 27 | 62.96 | 0.155 | 0.037 | 30.919 | 4.371 |
| Pycnonotidae | 15.44 | 118 | 151 | 78.15 | 0.035 | 0.012 | 19.706 | 4.054 |
| Rhinocryptidae | 17.46 | 29 | 65 | 44.62 | 0.029 | 0.028 | 14.802 | 3.122 |
| Rhipiduridae | 13.73 | 8 | 54 | 14.81 | 0.014 | 0.006 | 14.935 | 2.568 |
| Scotocercidae | 11.34 | 31 | 36 | 86.11 | 0.01 | 0.005 | 13.631 | 2.852 |
| Sittidae | 15.34 | 23 | 28 | 82.14 | 0.017 | 0.007 | 17.807 | 3.585 |
| Sturnidae | 12.07 | 108 | 123 | 87.8 | 0.09 | 0.048 | 25.055 | 4.658 |
| Sylviidae | 13.7 | 55 | 69 | 79.71 | 0.017 | 0.011 | 12.936 | 3.223 |
| Thamnophilidae | 15.54 | 159 | 237 | 67.09 | 0.021 | 0.015 | 18.394 | 3.661 |
| Thraupidae | 10.75 | 329 | 381 | 86.35 | 0.023 | 0.015 | 14.989 | 3.545 |
| Timaliidae | 11.43 | 52 | 54 | 96.3 | 0.027 | 0.019 | 20.928 | 8.91 |
| Tityridae | 13.86 | 28 | 33 | 84.85 | 0.03 | 0.016 | 17.173 | 4.571 |
| Troglodytidae | 15.02 | 69 | 86 | 80.23 | 0.02 | 0.009 | 19.362 | 4.324 |
| Turdidae | 15.32 | 140 | 175 | 80 | 0.069 | 0.027 | 22.939 | 4.4 |
| Tyrannidae | 17.68 | 296 | 425 | 69.65 | 0.018 | 0.014 | 15.581 | 5.161 |
| Vangidae | 12.24 | 29 | 39 | 74.36 | 0.036 | 0.024 | 22.99 | 10.133 |
| Viduidae | 10.93 | 18 | 20 | 90 | 0.015 | 0.003 | 10.63 | 0.991 |
| Vireonidae | 15.29 | 52 | 63 | 82.54 | 0.016 | 0.009 | 15.324 | 2.542 |
| Zosteropidae | 11.82 | 98 | 142 | 69.01 | 0.013 | 0.005 | 14.355 | 2.358 |

**Table S5.**
Summary statistics of ordinary least squares (OLS) allometric correction per subclade. SS: Sum of Squares, $R^2$: Pearson's R squared, F: F-statistic, Z: Z-score from RRPP permutations, Pr(>F): p-value.

| | SS | $R^2$ | F | Z | Pr(>F) |
|---|---|---|---|---|---|
| Paleognathae | 0.028849 | 0.026531 | 1.335448 | 0.689847 | 0.254975 |
| Anseriformes | 0.018384 | 0.00559 | 0.854479 | 0.12546 | 0.454855 |
| Galliformes | 0.193011 | 0.028844 | 7.751993 | 2.77279 | 0.0012 |
| Podicipediformes | 0.012357 | 0.062831 | 1.072701 | 0.477375 | 0.323568 |
| Columbiformes | 0.011761 | 0.003146 | 0.830031 | 0.162331 | 0.442456 |
| Pterocliformes | 0.002191 | 0.020941 | 0.299447 | -1.42119 | 0.924608 |
| Otidiformes | 0.042981 | 0.161942 | 4.444386 | 2.188616 | 0.010899 |
| Musophagiformes | 0.041895 | 0.048696 | 1.023763 | 0.557815 | 0.312969 |
| Cuculiformes | 0.062758 | 0.02974 | 3.892712 | 2.272003 | 0.008699 |
| Caprimulgiformes | 0.037396 | 0.001118 | 0.408652 | -0.16712 | 0.558044 |
| Gruiformes | 0.097473 | 0.035573 | 4.942639 | 2.338836 | 0.005699 |
| Charadriiformes | 0.25415 | 0.019222 | 6.898794 | 2.62625 | 0.0024 |
| Sphenisciformes | 0.006843 | 0.025449 | 0.391703 | -0.6406 | 0.733227 |
| Procellariiformes | 0.134605 | 0.083044 | 9.690423 | 3.742383 | 1.00E-04 |
| Ciconiiformes | 0.004795 | 0.086691 | 1.613638 | 0.942156 | 0.174883 |
| Suliformes | 0.013244 | 0.020081 | 1.024633 | 0.449501 | 0.333767 |
| Pelecaniformes | 0.059237 | 0.026061 | 2.67579 | 1.536998 | 0.064294 |
| Accipitriformes | 0.070113 | 0.019791 | 4.542791 | 2.461931 | 0.005399 |
| Strigiformes | 0.011974 | 0.007059 | 0.490561 | -0.34035 | 0.619838 |
| Trogoniformes | 0.001305 | 0.003503 | 0.13359 | -2.06146 | 0.982502 |
| Bucerotiformes | 0.052095 | 0.028602 | 1.854978 | 1.119183 | 0.139686 |
| Coraciiformes | 0.033118 | 0.010263 | 1.524359 | 0.883936 | 0.191681 |
| Galbuliformes | 0.087602 | 0.083355 | 3.728327 | 1.779127 | 0.037096 |
| Piciformes | 0.200598 | 0.025597 | 8.879084 | 3.041023 | 0.0004 |
| Falconiformes | 0.00923 | 0.010983 | 0.655216 | -0.25934 | 0.59954 |
| Psittaciformes | 0.018301 | 0.003654 | 1.239434 | 0.629571 | 0.269373 |
| Pittidae | 0.022006 | 0.121355 | 4.143488 | 2.092757 | 0.015798 |
| Thamnophilidae | 0.01344 | 0.010213 | 1.619991 | 0.969263 | 0.172583 |
| Grallariidae | 0.006973 | 0.024421 | 0.650841 | -0.13922 | 0.548845 |
| Rhinocryptidae | 0.005965 | 0.010223 | 0.278879 | -0.80237 | 0.778622 |
| Furnariidae | 0.050675 | 0.018196 | 4.392316 | 2.269375 | 0.008899 |
| Pipridae | 0.004166 | 0.006759 | 0.313027 | -0.73144 | 0.765523 |
| Cotingidae | 0.014253 | 0.014591 | 0.814376 | 0.189964 | 0.421658 |
| Tityridae | 0.006659 | 0.022516 | 0.598897 | -0.2841 | 0.606439 |
| Tyrannidae | 0.275383 | 0.065822 | 20.71502 | 4.86066 | 1.00E-04 |
| Ptilonorhynchidae | 0.01485 | 0.057431 | 0.913955 | 0.222595 | 0.416258 |
| Maluridae | 0.012921 | 0.07155 | 1.464217 | 0.79828 | 0.219878 |
| Meliphagidae | 0.029442 | 0.015583 | 2.422005 | 1.371068 | 0.090491 |
| Acanthizidae | 0.001295 | 0.002301 | 0.119935 | -2.02065 | 0.982802 |

| | | | | | |
|---|---|---|---|---|---|
| Campephagidae | 0.17369 | 0.19705 | 17.42395 | 3.553309 | 1.00E-04 |
| Vireonidae | 0.007514 | 0.012562 | 0.636095 | -0.14911 | 0.560444 |
| Pachycephalidae | 0.005321 | 0.016916 | 0.585037 | -0.13941 | 0.551045 |
| Oriolidae | 0.007368 | 0.024436 | 0.75143 | 0.067245 | 0.473853 |
| Artamidae | 0.08684 | 0.203779 | 5.118642 | 1.980932 | 0.019498 |
| Platysteiridae | 0.002758 | 0.026887 | 0.607849 | -0.25577 | 0.591541 |
| Vangidae | 0.011793 | 0.017384 | 0.477667 | -0.45799 | 0.670833 |
| Malaconotidae | 0.016046 | 0.047136 | 2.077626 | 1.215708 | 0.120988 |
| Rhipiduridae | 0.010931 | 0.11128 | 0.75128 | -0.08522 | 0.552045 |
| Dicruridae | 0.008397 | 0.033456 | 0.519204 | -0.14549 | 0.544346 |
| Paradisaeidae | 0.068719 | 0.041187 | 1.589389 | 0.887761 | 0.19718 |
| Monarchidae | 0.042489 | 0.133794 | 3.707045 | 1.984604 | 0.023098 |
| Laniidae | 0.004455 | 0.025611 | 0.762252 | -0.14347 | 0.558044 |
| Corvidae | 0.052488 | 0.026145 | 2.899476 | 1.666129 | 0.048795 |
| Petroicidae | 0.008779 | 0.028173 | 0.956654 | 0.297573 | 0.393461 |
| Paridae | 0.008543 | 0.014204 | 0.749243 | 0.056204 | 0.474153 |
| Alaudidae | 0.020155 | 0.016387 | 1.399397 | 0.761511 | 0.232377 |
| Macrosphenidae | 0.004515 | 0.033324 | 0.517098 | -0.51352 | 0.694031 |
| Acrocephalidae | 0.012665 | 0.033424 | 1.659841 | 0.987274 | 0.166283 |
| Cisticolidae | 0.001554 | 0.001764 | 0.240314 | -1.36116 | 0.910509 |
| Locustellidae | 0.008845 | 0.025434 | 1.148294 | 0.522279 | 0.30067 |
| Hirundinidae | 0.009727 | 0.016868 | 1.269649 | 0.675318 | 0.250375 |
| Pycnonotidae | 0.036031 | 0.033977 | 4.079999 | 2.153348 | 0.012299 |
| Phylloscopidae | 0.016416 | 0.038051 | 2.452477 | 1.428176 | 0.081192 |
| Scotocercidae | 0.01765 | 0.063227 | 1.957355 | 1.211164 | 0.115488 |
| Sylviidae | 0.044306 | 0.016996 | 0.916354 | 0.411758 | 0.351065 |
| Zosteropidae | 0.001553 | 0.002427 | 0.23353 | -1.31963 | 0.90411 |
| Timaliidae | 0.007943 | 0.010538 | 0.532514 | -0.31026 | 0.616238 |
| Pellorneidae | 0.026869 | 0.042081 | 2.284326 | 1.309769 | 0.10039 |
| Leiothrichidae | 0.010755 | 0.009353 | 1.217887 | 0.607692 | 0.275372 |
| Sittidae | 0.00791 | 0.034836 | 0.757969 | 0.140134 | 0.439856 |
| Polioptilidae | 0.027474 | 0.495834 | 7.86779 | 2.35205 | 0.0048 |
| Troglodytidae | 0.004163 | 0.008239 | 0.556607 | -0.31437 | 0.615938 |
| Sturnidae | 0.008037 | 0.00469 | 0.49948 | -0.41522 | 0.652735 |
| Mimidae | 0.01163 | 0.041618 | 1.389621 | 0.735332 | 0.241576 |
| Turdidae | 0.032197 | 0.020601 | 2.902741 | 1.684444 | 0.045595 |
| Muscicapidae | 0.008991 | 0.00277 | 0.775024 | 0.081184 | 0.470853 |
| Dicaeidae | 0.028396 | 0.056239 | 2.383591 | 1.441156 | 0.076092 |
| Nectariniidae | 0.00934 | 0.011821 | 1.447409 | 0.821264 | 0.212279 |
| Ploceidae | 0.041585 | 0.033379 | 3.729472 | 1.908961 | 0.024098 |
| Estrildidae | 0.016803 | 0.010763 | 1.39265 | 0.787966 | 0.225277 |
| Viduidae | 0.001775 | 0.014934 | 0.242561 | -1.14869 | 0.870713 |
| Passeridae | 0.005256 | 0.017919 | 0.69334 | -0.22517 | 0.581042 |
| Motacillidae | 0.007549 | 0.015797 | 0.914886 | 0.300592 | 0.389461 |
| Fringillidae | 0.074814 | 0.014954 | 2.808541 | 1.663999 | 0.047195 |

| | | | | | |
|---|---|---|---|---|---|
| Emberizidae | 0.002369 | 0.006191 | 0.242935 | -1.57571 | 0.944706 |
| Passerellidae | 0.002315 | 0.002276 | 0.244046 | -1.4311 | 0.925407 |
| Icteridae | 0.122844 | 0.093557 | 9.495618 | 3.001247 | 0.0003 |
| Parulidae | 0.031013 | 0.036754 | 3.319592 | 1.789604 | 0.034297 |
| Cardinalidae | 0.021395 | 0.03819 | 1.707354 | 1.064303 | 0.149785 |
| Thraupidae | 0.026005 | 0.003178 | 1.042425 | 0.398811 | 0.361064 |

**Table S6.**
Pagel's λ per subclade, estimated via fit_t_pl() in RPANDA v2.5 for each residual landmark set per subclade. Residual landmarks calculated from the multivariate regression of landmarks onto (log) centroid size using procD.lm() in geomorph v4.0.1.

| | λ |
|---|---|
| Paleognathae | 0.672558 |
| Anseriformes | 0.503307 |
| Galliformes | 0.047413 |
| Podicipediformes | 0.336736 |
| Columbiformes | 0.131534 |
| Pterocliformes | 0.253068 |
| Otidiformes | 0.160588 |
| Musophagiformes | 0.415798 |
| Cuculiformes | 0.159896 |
| Caprimulgiformes | 0.913667 |
| Gruiformes | 0.148168 |
| Charadriiformes | 0.160545 |
| Sphenisciformes | 0.70182 |
| Procellariiformes | 0.713587 |
| Ciconiiformes | 0.546153 |
| Suliformes | 0.674546 |
| Pelecaniformes | 0.764723 |
| Accipitriformes | 0.187068 |
| Strigiformes | 0.086126 |
| Trogoniformes | 0.255638 |
| Bucerotiformes | 0.540625 |
| Coraciiformes | 0.372411 |
| Galbuliformes | 0.375492 |
| Piciformes | 0.137949 |
| Falconiformes | 0.462688 |
| Psittaciformes | 0.275641 |
| Pittidae | 0.132526 |
| Thamnophilidae | 0.104108 |
| Grallariidae | 1.00E-05 |
| Rhinocryptidae | 0.087097 |
| Furnariidae | 0.102482 |
| Pipridae | 0.230572 |
| Cotingidae | 0.321181 |
| Tityridae | 0.516398 |
| Tyrannidae | 0.183237 |
| Ptilonorhynchidae | 0.406739 |
| Maluridae | 0.342937 |
| Meliphagidae | 0.036389 |
| Acanthizidae | 0.360257 |

| | |
|---|---|
| Campephagidae | 0.141808 |
| Vireonidae | 0.425348 |
| Pachycephalidae | 0.31293 |
| Oriolidae | 0.340218 |
| Artamidae | 1.00E-05 |
| Platysteiridae | 0.075319 |
| Vangidae | 0.292372 |
| Malaconotidae | 0.217804 |
| Rhipiduridae | 0.937543 |
| Dicruridae | 0.321077 |
| Paradisaeidae | 0.344192 |
| Monarchidae | 0.07871 |
| Laniidae | 0.385813 |
| Corvidae | 0.185587 |
| Petroicidae | 0.07708 |
| Paridae | 0.099065 |
| Alaudidae | 0.095853 |
| Macrosphenidae | 0.310593 |
| Acrocephalidae | 0.257152 |
| Cisticolidae | 0.138749 |
| Locustellidae | 0.10742 |
| Hirundinidae | 0.155588 |
| Pycnonotidae | 0.213083 |
| Phylloscopidae | 0.112573 |
| Scotocercidae | 0.214796 |
| Sylviidae | 0.242034 |
| Zosteropidae | 0.080122 |
| Timaliidae | 0.17512 |
| Pellorneidae | 0.246094 |
| Leiothrichidae | 0.112988 |
| Sittidae | 0.12427 |
| Polioptilidae | 0.017078 |
| Troglodytidae | 0.173909 |
| Sturnidae | 0.168614 |
| Mimidae | 0.295507 |
| Turdidae | 0.104407 |
| Muscicapidae | 0.056927 |
| Dicaeidae | 0.330835 |
| Nectariniidae | 0.088574 |
| Ploceidae | 0.20532 |
| Estrildidae | 0.099705 |
| Viduidae | 0.237573 |
| Passeridae | 0.226946 |
| Motacillidae | 0.154549 |
| Fringillidae | 0.175791 |

| | |
|---|---|
| Emberizidae | 0.2417 |
| Passerellidae | 0.112166 |
| Icteridae | 0.111534 |
| Parulidae | 0.141545 |
| Cardinalidae | 0.301305 |
| Thraupidae | 0.066832 |

**Table S7.**
Summary statistics of phylogenetic generalized least squares (PGLS) allometric correction per subclade. SS: Sum of Squares, MS: mean squared error, $R^2$: Pearson's R squared, F: F-statistic, Z: Z-score from RRPP permutations, Pr(>F): p-value.

| | SS | $R^2$ | F | Z | Pr(>F) |
|---|---|---|---|---|---|
| Paleognathae | 0.201962 | 0.082869 | 4.427497 | 1.739681 | 0.037896 |
| Anseriformes | 0.014549 | 0.002754 | 0.419782 | -1.0951 | 0.857014 |
| Galliformes | 0.116339 | 0.018703 | 4.9744 | 2.235118 | 0.008699 |
| Podicipediformes | 0.043866 | 0.045428 | 0.761433 | 0.155312 | 0.439856 |
| Columbiformes | 0.0024 | 0.000728 | 0.19173 | -1.62065 | 0.947105 |
| Pterocliformes | 0.05947 | 0.100337 | 1.561375 | 0.862388 | 0.210379 |
| Otidiformes | 0.289478 | 0.288754 | 9.337609 | 2.758674 | 0.0015 |
| Musophagiformes | 0.083197 | 0.03924 | 0.816847 | 0.194936 | 0.433057 |
| Cuculiformes | 0.51784 | 0.073301 | 10.04555 | 2.587685 | 0.0017 |
| Caprimulgiformes | 0.010946 | 0.000854 | 0.31191 | -0.95461 | 0.826417 |
| Gruiformes | 0.070611 | 0.009917 | 1.342238 | 0.784559 | 0.216978 |
| Charadriiformes | 0.093144 | 0.013374 | 4.771621 | 2.337573 | 0.006799 |
| Sphenisciformes | 0.216493 | 0.183155 | 3.363337 | 1.456679 | 0.075692 |
| Procellariiformes | 0.035052 | 0.01249 | 1.353308 | 1.192427 | 0.121488 |
| Ciconiiformes | 0.131333 | 0.135381 | 2.661849 | 1.242685 | 0.115688 |
| Suliformes | 0.067838 | 0.033741 | 1.745941 | 0.691393 | 0.256074 |
| Pelecaniformes | 0.056423 | 0.014625 | 1.484164 | 0.843487 | 0.20378 |
| Accipitriformes | 0.054428 | 0.008393 | 1.904308 | 1.372448 | 0.085491 |
| Strigiformes | 0.271067 | 0.061039 | 4.485509 | 2.136768 | 0.013699 |
| Trogoniformes | 0.13752 | 0.054092 | 2.173022 | 1.09216 | 0.152585 |
| Bucerotiformes | 0.16465 | 0.051538 | 3.423359 | 1.244025 | 0.109689 |
| Coraciiformes | 0.183188 | 0.032528 | 4.94239 | 1.494004 | 0.068193 |
| Galbuliformes | 0.276412 | 0.105407 | 4.830919 | 1.849138 | 0.029497 |
| Piciformes | 0.116889 | 0.028552 | 9.934043 | 3.356332 | 1.00E-04 |
| Falconiformes | 0.083695 | 0.031819 | 1.939045 | 0.830677 | 0.217578 |
| Psittaciformes | 0.017138 | 0.003939 | 1.336477 | 0.668295 | 0.260574 |
| Pittidae | 0.035368 | 0.037445 | 1.167041 | 0.588594 | 0.289071 |
| Thamnophilidae | 0.171044 | 0.025319 | 4.078396 | 1.584208 | 0.056494 |
| Grallariidae | 0.006973 | 0.02442 | 0.650826 | -0.13924 | 0.548845 |
| Rhinocryptidae | 0.137845 | 0.126784 | 3.920177 | 2.101878 | 0.015098 |
| Furnariidae | 0.018268 | 0.007254 | 1.73176 | 1.172871 | 0.126587 |
| Pipridae | 0.076993 | 0.030125 | 1.428771 | 0.755268 | 0.236576 |
| Cotingidae | 0.09264 | 0.026397 | 1.491195 | 0.784523 | 0.222778 |
| Tityridae | 0.015223 | 0.008389 | 0.219959 | -0.82396 | 0.779022 |
| Tyrannidae | 0.101633 | 0.027273 | 8.243045 | 3.298504 | 1.00E-04 |
| Ptilonorhynchidae | 0.044683 | 0.046244 | 0.727292 | -0.02797 | 0.508549 |
| Maluridae | 0.08918 | 0.086436 | 1.797675 | 0.988077 | 0.174583 |
| Meliphagidae | 0.103065 | 0.013972 | 2.168057 | 1.221202 | 0.117488 |
| Acanthizidae | 0.314325 | 0.113632 | 6.666372 | 2.042698 | 0.014399 |

| | | | | | |
|---|---|---|---|---|---|
| Campephagidae | 0.812962 | 0.211411 | 19.03423 | 2.967708 | 0.0002 |
| Vireonidae | 0.012609 | 0.004385 | 0.220221 | -0.94235 | 0.819118 |
| Pachycephalidae | 0.089227 | 0.042585 | 1.512308 | 0.748341 | 0.240376 |
| Oriolidae | 0.013684 | 0.007653 | 0.231367 | -0.78284 | 0.775022 |
| Artamidae | 0.086833 | 0.203767 | 5.118285 | 1.980877 | 0.019498 |
| Platysteiridae | 0.00821 | 0.024503 | 0.552599 | -0.20274 | 0.571743 |
| Vangidae | 0.158393 | 0.097573 | 2.919305 | 1.673293 | 0.043096 |
| Malaconotidae | 0.02844 | 0.014332 | 0.6107 | 0.046752 | 0.482552 |
| Rhipiduridae | 0.066101 | 0.097232 | 0.646229 | -0.06197 | 0.507949 |
| Dicruridae | 0.084115 | 0.080225 | 1.308328 | 0.633091 | 0.270273 |
| Paradisaeidae | 0.042932 | 0.013283 | 0.498084 | -0.41876 | 0.662034 |
| Monarchidae | 0.112872 | 0.163087 | 4.676816 | 2.285196 | 0.010299 |
| Laniidae | 0.181076 | 0.100642 | 3.245228 | 1.349621 | 0.094391 |
| Corvidae | 0.49768 | 0.087964 | 10.41632 | 2.704093 | 0.0009 |
| Petroicidae | 0.018378 | 0.034352 | 1.173943 | 0.510867 | 0.309869 |
| Paridae | 0.101343 | 0.047993 | 2.621453 | 1.413459 | 0.082692 |
| Alaudidae | 0.017327 | 0.004993 | 0.421485 | -0.51753 | 0.689731 |
| Macrosphenidae | 0.01798 | 0.028648 | 0.442392 | -0.38356 | 0.643936 |
| Acrocephalidae | 0.54016 | 0.256313 | 16.54325 | 2.853344 | 0.0002 |
| Cisticolidae | 0.066535 | 0.012374 | 1.703885 | 0.951109 | 0.191281 |
| Locustellidae | 0.029326 | 0.025074 | 1.131623 | 0.543813 | 0.29827 |
| Hirundinidae | 0.098299 | 0.034399 | 2.636211 | 1.415116 | 0.081392 |
| Pycnonotidae | 0.061237 | 0.010649 | 1.248582 | 0.688495 | 0.265273 |
| Phylloscopidae | 0.186649 | 0.065487 | 4.34472 | 1.795694 | 0.031897 |
| Scotocercidae | 0.166741 | 0.091422 | 2.918013 | 1.406432 | 0.084392 |
| Sylviidae | 0.028214 | 0.007687 | 0.410543 | -0.62066 | 0.725527 |
| Zosteropidae | 0.019665 | 0.004044 | 0.389764 | -0.27919 | 0.59654 |
| Timaliidae | 0.041434 | 0.022238 | 1.137177 | 0.505133 | 0.309369 |
| Pellorneidae | 0.07556 | 0.028191 | 1.508464 | 0.853066 | 0.205679 |
| Leiothrichidae | 0.90714 | 0.150291 | 22.81666 | 3.237071 | 1.00E-04 |
| Sittidae | 0.015361 | 0.025525 | 0.550067 | -0.24209 | 0.59644 |
| Polioptilidae | 0.027024 | 0.418782 | 5.764208 | 2.367825 | 0.008399 |
| Troglodytidae | 0.039061 | 0.015433 | 1.050236 | 0.516861 | 0.310869 |
| Sturnidae | 0.208658 | 0.039783 | 4.391717 | 2.044296 | 0.019198 |
| Mimidae | 0.096007 | 0.065944 | 2.25919 | 1.232069 | 0.117588 |
| Turdidae | 0.67548 | 0.073215 | 10.90192 | 2.396872 | 0.0018 |
| Muscicapidae | 0.013061 | 0.004445 | 1.24574 | 0.638087 | 0.270473 |
| Dicaeidae | 0.098379 | 0.032712 | 1.352724 | 0.728944 | 0.248575 |
| Nectariniidae | 0.07744 | 0.012298 | 1.506638 | 0.845841 | 0.217078 |
| Ploceidae | 0.234373 | 0.048514 | 5.506627 | 1.718215 | 0.039296 |
| Estrildidae | 0.068012 | 0.012364 | 1.602374 | 0.94997 | 0.187681 |
| Viduidae | 0.283704 | 0.269273 | 5.895992 | 1.953712 | 0.017198 |
| Passeridae | 0.007595 | 0.003125 | 0.119122 | -1.29761 | 0.89781 |
| Motacillidae | 0.171721 | 0.084082 | 5.232671 | 2.129352 | 0.011999 |
| Fringillidae | 0.128371 | 0.011931 | 2.233812 | 1.237172 | 0.114089 |

| | | | | | |
|---|---|---|---|---|---|
| Emberizidae | 0.015916 | 0.008497 | 0.334206 | -0.60361 | 0.712829 |
| Passerellidae | 0.170066 | 0.033284 | 3.684001 | 1.715229 | 0.041096 |
| Icteridae | 0.280678 | 0.054605 | 5.313838 | 2.087046 | 0.014699 |
| Parulidae | 0.220892 | 0.055162 | 5.079315 | 1.883685 | 0.025797 |
| Cardinalidae | 0.364919 | 0.184302 | 9.715615 | 2.73106 | 0.0011 |
| Thraupidae | 0.015759 | 0.002434 | 0.797765 | 0.07085 | 0.476252 |

**Table S8.**
$AIC_c$ comparisons between OLS and PGLS regressions for allometric corrections.

| | ols | pgls | best fitting model |
|---|---|---|---|
| Paleognathae | -3391.82 | -4608.43 | pgls |
| Anseriformes | -33368.8 | -45180.9 | pgls |
| Galliformes | -95264 | -96479.6 | pgls |
| Podicipediformes | -353.458 | -382.483 | pgls |
| Columbiformes | -96651.4 | -99847 | pgls |
| Pterocliformes | -273.267 | -281.781 | pgls |
| Otidiformes | -746.022 | -761.201 | pgls |
| Musophagiformes | -531.193 | -579.428 | pgls |
| Cuculiformes | -23314.1 | -24086.8 | pgls |
| Caprimulgiformes | -155122 | -276814 | pgls |
| Gruiformes | -25954.4 | -27185.8 | pgls |
| Charadriiformes | -147828 | -154735 | pgls |
| Sphenisciformes | -299.807 | -406.336 | pgls |
| Procellariiformes | -16534.5 | -24129.9 | pgls |
| Ciconiiformes | -418.235 | -467.138 | pgls |
| Suliformes | -3582.12 | -5158.81 | pgls |
| Pelecaniformes | -14387.5 | -20580.8 | pgls |
| Accipitriformes | -73516.3 | -79396.5 | pgls |
| Strigiformes | -6790.43 | -6891.55 | pgls |
| Trogoniformes | -2066.82 | -2211.43 | pgls |
| Bucerotiformes | -5623.33 | -7150.51 | pgls |
| Coraciiformes | -31227.7 | -36796.9 | pgls |
| Galbuliformes | -2365.81 | -2653.78 | pgls |
| Piciformes | -140040 | -146209 | pgls |
| Falconiformes | -4981.29 | -6303.88 | pgls |
| Psittaciformes | -140131 | -154598 | pgls |
| Pittidae | -1297.76 | -1319.79 | pgls |
| Thamnophilidae | -35851.3 | -36610.5 | pgls |
| Grallariidae | -955.355 | -955.355 | ols |
| Rhinocryptidae | -1011.58 | -1016.81 | pgls |
| Furnariidae | -81613.2 | -83691.9 | pgls |
| Pipridae | -3028.11 | -3209.88 | pgls |
| Cotingidae | -4317.11 | -4676.1 | pgls |
| Tityridae | -952.535 | -1087.72 | pgls |
| Tyrannidae | -114683 | -120028 | pgls |
| Ptilonorhynchidae | -300.28 | -330.178 | pgls |
| Maluridae | -506.552 | -552.602 | pgls |
| Meliphagidae | -33994.2 | -34259 | pgls |
| Acanthizidae | -3883.78 | -4234.21 | pgls |
| Campephagidae | -7287.28 | -7524.4 | pgls |

| | | | |
|---|---|---|---|
| Vireonidae | -3584.13 | -4037.93 | pgls |
| Pachycephalidae | -1655.5 | -1844.71 | pgls |
| Oriolidae | -1281.5 | -1384.2 | pgls |
| Artamidae | -549.327 | -549.327 | ols |
| Platysteiridae | -696.744 | -700.019 | pgls |
| Vangidae | -993.977 | -1016.72 | pgls |
| Malaconotidae | -2542.69 | -2617.47 | pgls |
| Rhipiduridae | -41.9947 | -50.2267 | pgls |
| Dicruridae | -305.109 | -322.879 | pgls |
| Paradisaeidae | -1884.72 | -2083.62 | pgls |
| Monarchidae | -808.068 | -813.358 | pgls |
| Laniidae | -1208.33 | -1355.08 | pgls |
| Corvidae | -16827.1 | -17520.1 | pgls |
| Petroicidae | -1555.2 | -1560.34 | pgls |
| Paridae | -3882.12 | -3926.22 | pgls |
| Alaudidae | -10170.8 | -10282.6 | pgls |
| Macrosphenidae | -312.283 | -318.961 | pgls |
| Acrocephalidae | -3322.77 | -3552.09 | pgls |
| Cisticolidae | -26916.2 | -27512.7 | pgls |
| Locustellidae | -2788.81 | -2815.28 | pgls |
| Hirundinidae | -7930.56 | -8071.19 | pgls |
| Pycnonotidae | -19537.2 | -20106 | pgls |
| Phylloscopidae | -5570.36 | -5720.98 | pgls |
| Scotocercidae | -1196.43 | -1263.87 | pgls |
| Sylviidae | -3928.88 | -4097.3 | pgls |
| Zosteropidae | -13392.2 | -13705.7 | pgls |
| Timaliidae | -3570.48 | -3653.06 | pgls |
| Pellorneidae | -3883.94 | -3996.77 | pgls |
| Leiothrichidae | -24173.3 | -24595.6 | pgls |
| Sittidae | -620.902 | -625.756 | pgls |
| Polioptilidae | -90.4527 | -90.4701 | pgls |
| Troglodytidae | -6501.2 | -6620.37 | pgls |
| Sturnidae | -16223.6 | -16645.3 | pgls |
| Mimidae | -1464.1 | -1542.12 | pgls |
| Turdidae | -27643.3 | -28209.5 | pgls |
| Muscicapidae | -106041 | -106493 | pgls |
| Dicaeidae | -2280.59 | -2432.82 | pgls |
| Nectariniidae | -21304.4 | -21552.1 | pgls |
| Ploceidae | -16903.4 | -17899.5 | pgls |
| Estrildidae | -23740.5 | -24084.6 | pgls |
| Viduidae | -361.186 | -388.094 | pgls |
| Passeridae | -2073.98 | -2161.91 | pgls |
| Motacillidae | -4694.54 | -4771.92 | pgls |
| Fringillidae | -49472.3 | -51503.4 | pgls |
| Emberizidae | -2174.99 | -2244.77 | pgls |

| | | | |
|---|---|---|---|
| Passerellidae | -16603.2 | -16838.8 | pgls |
| Icteridae | -12232.7 | -12455.3 | pgls |
| Parulidae | -10964.5 | -11261.9 | pgls |
| Cardinalidae | -2633.62 | -2736.33 | pgls |
| Thraupidae | -133538 | -134280 | pgls |

**Table S9.**
PGLS of simple and SCM-based GCs on (log) species richness and clade age per subclade.

| **PGLS MODEL** | $\mathbf{R^2}$ | **P** | **F** |
|---|---|---|---|
| Species Richness (SR) | | | |
| Simple Gini ~ SR | 0.040 | 0.059 | 3.674 |
| SCM Gini ~ SR | 0.034 | 0.080 | 3.136 |
| Clade Age (CA) | | | |
| Simple Gini ~ Clade Age | 0.000 | 0.862 | 0.030 |
| SCM Gini ~ Clade Age | 0.001 | 0.757 | 0.097 |

**Table S10.** All four macroevolutionary variables regressed onto clade age, species richness, or an interaction between clade age and species richness. Preferred models according to $AIC_c$ values are shown in bold.

| PGLS Model | $R^2$ | p | F | Z | AICc | ΔAICc |
|---|---|---|---|---|---|---|
| Integration ~ Clade Age | **0.014** | **0.305** | **1.102** | **0.313** | **-12.316** | **0** |
| Integration ~ Species Richness | 0.002 | 0.715 | 0.140 | -0.781 | -11.337 | 0.978 |
| Integration ~ Clade Age*Species Richness | -0.013 | 0.641 | 0.563 | -0.479 | -8.951 | 3.365 |
| Rates ~ Clade Age | **0.101** | **0.003** | **9.052** | **2.483** | **150.477** | **0** |
| Rates ~ Species Richness | 0.015 | 0.270 | 1.235 | 1.917 | 156.879 | 6.402 |
| Rates ~ Clade Age*Species Richness | 0.111 | 0.007 | 4.333 | 2.650 | 150.626 | 0.149 |
| Disparity ~ Clade Age | 0.226 | 0.000 | 23.099 | 3.340 | 92.389 | 10.970 |
| Disparity ~ Species Richness | **0.150** | **0.000** | **13.989** | **7.105** | **81.419** | **0** |
| Disparity ~ Clade Age*Species Richness | 0.302 | 0.000 | 12.403 | 4.274 | 85.233 | 3.814 |
| Ecological Specialization ~ Clade Age | 0.021 | 0.192 | 1.730 | 0.920 | 396.988 | 6.716 |
| Ecological Specialization ~ Species Richness | **0.097** | **0.005** | **8.703** | **2.317** | **390.272** | **0** |
| Ecological Specialization ~ Clade Age*Species Richness | 0.069 | 0.038 | 2.941 | 1.762 | 393.955 | 3.683 |

**Table S11.** Per-subclade (90) results for disparity (Procrustes Variance), rates (Brownian motion of λ-rescaled branches), integration (Z-score), and ecological specialization (Gini coefficient). Italicized subclades are Passeriforme families.

| Clade | Disparity | Rates | Integration | Gini |
|---|---|---|---|---|
| Paleognathae | 0.0240714 | 2.21E-06 | -0.19417115 | 0.922764623 |
| Anseriformes | 0.0307545 | 2.51E-05 | -0.38945179 | 0.32433543 |
| Galliformes | 0.0253036 | 1.94E-05 | 0.192241907 | 0.623900485 |
| Podicipediformes | 0.0108918 | 3.95E-06 | -0.25076326 | 1 |
| Columbiformes | 0.0142208 | 6.47E-06 | -0.0817046 | 0.25649844 |
| Pterocliformes | 0.0064165 | 2.40E-06 | -0.59549796 | 0.851132152 |
| Otidiformes | 0.0089587 | 1.07E-05 | -0.20231926 | 0.846067613 |
| Musophagiformes | 0.0386393 | 5.68E-06 | 0.640215552 | 0.946890951 |
| Cuculiformes | 0.0163151 | 2.87E-06 | -0.35121245 | 0.780105453 |
| Caprimulgiformes | 0.3296536 | 5.98E-06 | 0.027078555 | 0.757891558 |
| Gruiformes | 0.0205474 | 5.09E-06 | -0.16716926 | 0.552206362 |
| Charadriiformes | 0.0371905 | 7.33E-06 | -0.36600093 | 0.614772577 |
| Sphenisciformes | 0.0191745 | 6.02E-06 | -0.24284926 | 1 |
| Procellariiformes | 0.0169162 | 3.13E-06 | -0.61678259 | 0.98845874 |
| Ciconiiformes | 0.0027239 | 1.39E-06 | 0.11754502 | 0.778215809 |
| Suliformes | 0.0176903 | 6.45E-06 | 0.129890604 | 1 |
| Pelecaniformes | 0.0243072 | 2.29E-06 | -0.18505012 | 0.575345261 |
| Accipitriformes | 0.0160181 | 1.12E-05 | -0.04868537 | 0.836726806 |
| Strigiformes | 0.023838 | 9.94E-06 | 0.140249116 | 0.58619264 |
| Trogoniformes | 0.009438 | 5.45E-06 | -0.39247523 | 0.313144967 |
| Bucerotiformes | 0.0374106 | 4.92E-06 | -0.3975887 | 0.434839474 |
| Coraciiformes | 0.0230556 | 4.81E-06 | -0.43490911 | 0.721480175 |
| Galbuliformes | 0.0247486 | 4.18E-06 | 0.037731818 | 1 |
| Piciformes | 0.0283952 | 7.33E-06 | -0.15641672 | 0.682763552 |
| Falconiformes | 0.0154047 | 1.85E-05 | -0.08832091 | 0.412438379 |
| Psittaciformes | 0.0183146 | 8.22E-06 | -0.66218623 | 0.530555378 |
| *Pittidae* | 0.0049928 | 6.08E-06 | 0.21914254 | 1 |
| *Thamnophilidae* | 0.0082996 | 4.81E-06 | 0.041929082 | 1 |
| *Grallariidae* | 0.0099492 | 4.98E-06 | 0.135943179 | 0.97033763 |
| *Rhinocryptidae* | 0.0199294 | 6.46E-06 | 0.248553445 | 1 |
| *Furnariidae* | 0.0117095 | 8.23E-06 | -0.09699023 | 0.99700091 |
| *Pipridae* | 0.0131172 | 1.56E-05 | 0.174778191 | 0.815247856 |
| *Cotingidae* | 0.0171748 | 2.09E-05 | -0.19684751 | 0.895356272 |
| *Tityridae* | 0.0107426 | 5.08E-06 | -0.19783486 | 0.347082574 |
| *Tyrannidae* | 0.0145914 | 1.10E-05 | -0.10546873 | 0.969030562 |
| *Ptilonorhynchidae* | 0.0145279 | 1.25E-05 | 0.250980355 | 0.204429165 |
| *Maluridae* | 0.0081825 | 4.65E-06 | -0.12509369 | 0.599311245 |
| *Meliphagidae* | 0.0120492 | 5.65E-06 | -0.12549687 | 0.529549507 |
| *Acanthizidae* | 0.0106874 | 8.30E-06 | -0.05909713 | 1 |
| *Campephagidae* | 0.010338 | 8.09E-06 | 0.058342455 | 0.883113183 |

| | | | | |
|---|---|---|---|---|
| *Vireonidae* | 0.0122533 | 5.54E-06 | -0.14870799 | 0.98035966 |
| *Pachycephalidae* | 0.0087755 | 1.28E-05 | 0.065103655 | 0.655650924 |
| *Oriolidae* | 0.0101769 | 5.85E-06 | -0.29722372 | 0.0152177 |
| *Artamidae* | 0.0154231 | 1.27E-05 | -0.19447216 | 0.464023174 |
| *Platysteiridae* | 0.0041696 | 2.96E-06 | -0.38444627 | 1 |
| *Vangidae* | 0.0232126 | 9.68E-06 | -0.239135 | 1 |
| *Malaconotidae* | 0.0074551 | 5.03E-06 | 0.13736713 | 0.951983346 |
| *Rhipiduridae* | 0.0115707 | 4.24E-06 | -0.19225898 | 1 |
| *Dicruridae* | 0.0147316 | 1.50E-05 | 0.531909772 | 1 |
| *Paradisaeidae* | 0.0425343 | 3.36E-05 | 0.551433131 | 0.748897339 |
| *Monarchidae* | 0.0106989 | 8.72E-06 | -0.25757427 | 0.901798673 |
| *Laniidae* | 0.0057651 | 6.88E-06 | -0.5969151 | 0.96477946 |
| *Corvidae* | 0.0187772 | 1.41E-05 | 0.005783117 | 0.859848759 |
| *Petroicidae* | 0.0086576 | 2.43E-06 | -0.42738056 | 1 |
| *Paridae* | 0.0109987 | 5.71E-06 | -0.30909405 | 0.956961539 |
| *Alaudidae* | 0.0141144 | 7.46E-06 | -0.33070101 | 0.501406284 |
| *Macrosphenidae* | 0.0077538 | 3.07E-06 | -0.3303019 | 1 |
| *Acrocephalidae* | 0.007656 | 9.64E-06 | -0.1238552 | 1 |
| *Cisticolidae* | 0.0065074 | 4.36E-06 | -0.14278878 | 1 |
| *Locustellidae* | 0.0073713 | 5.49E-06 | -0.20707352 | 1 |
| *Hirundinidae* | 0.0074783 | 6.81E-06 | -0.04886754 | 1 |
| *Pycnonotidae* | 0.0087105 | 4.72E-06 | -0.30737155 | 0.200454348 |
| *Phylloscopidae* | 0.0065121 | 0.000622 | 1.215036719 | 1 |
| *Scotocercidae* | 0.008923 | 6.05E-05 | -0.18122621 | 1 |
| *Sylviidae* | 0.0473984 | 1.38E-05 | -0.18201939 | 0.778184638 |
| *Zosteropidae* | 0.0067063 | 1.66E-05 | -0.24755321 | 0.546902574 |
| *Timaliidae* | 0.0146099 | 8.03E-06 | -0.22103013 | 1 |
| *Pellorneidae* | 0.0115528 | 5.01E-06 | -0.04024945 | 1 |
| *Leiothrichidae* | 0.0088044 | 9.70E-06 | -0.08047222 | 0.813776148 |
| *Sittidae* | 0.0095386 | 5.49E-06 | -0.09883224 | 0.739604794 |
| *Polioptilidae* | 0.0027943 | 3.16E-06 | -0.2954333 | 1 |
| *Troglodytidae* | 0.0072836 | 4.44E-06 | 0.08794301 | 1 |
| *Sturnidae* | 0.0159141 | 1.29E-05 | 0.073137041 | 0.364016573 |
| *Mimidae* | 0.0079402 | 6.02E-06 | -0.38906701 | 0.019290795 |
| *Turdidae* | 0.0114744 | 8.23E-06 | -0.01782373 | 0.50433302 |
| *Muscicapidae* | 0.0115646 | 5.13E-06 | -0.18656516 | 0.934013056 |
| *Dicaeidae* | 0.011854 | 4.22E-05 | 0.157374434 | 0.620364368 |
| *Nectariniidae* | 0.0064269 | 3.47E-06 | -0.21421037 | 0.436180477 |
| *Ploceidae* | 0.0117191 | 1.29E-05 | -0.21781952 | 0.291129383 |
| *Estrildidae* | 0.0119512 | 1.98E-05 | -0.60616332 | 0.896488205 |
| *Viduidae* | 0.0067658 | 2.68E-05 | 0.143823895 | 1 |
| *Passeridae* | 0.0074499 | 8.07E-06 | -0.19444848 | 0.916623564 |
| *Motacillidae* | 0.0079961 | 5.03E-06 | -0.09044668 | 0.96865162 |
| *Fringillidae* | 0.0268238 | 1.64E-05 | -0.46553362 | 0.5547517 |
| *Emberizidae* | 0.0092976 | 7.88E-06 | -0.37793454 | 0.017080796 |

| | | | | |
|---|---|---|---|---|
| *Passerellidae* | 0.0093361 | 1.09E-05 | -0.25470954 | 0.718562127 |
| *Icteridae* | 0.0130618 | 1.68E-05 | -0.01949075 | 0.674210379 |
| *Parulidae* | 0.0092 | 1.22E-05 | -0.19906003 | 0.981932683 |
| *Cardinalidae* | 0.0119982 | 7.36E-06 | -0.55706789 | 0.419316599 |
| *Thraupidae* | 0.0248897 | 2.58E-05 | -0.41275683 | 0.439455265 |
